# Glycoprotein G enables HSV-2 neuroinvasion and provides protection as a glycosylated vaccine antigen

**DOI:** 10.64898/2026.02.25.706656

**Authors:** Ebba Könighofer, Carolina Gustafsson, Lindvi Gudmundsdotter, Ekaterina Mirgorodskaya, Jonas Nilsson, Maria Ekblad, Beata Adamiak, Eva Jennische, Stefan Lange, Edward Trybala, Staffan Görander, Tomas Bergström, Jan-Åke Liljeqvist, Rickard Nordén

**Affiliations:** Department of Infectious Diseases, Institute of Biomedicine, Sahlgrenska Academy, University of Gothenburg, Gothenburg, Sweden; Department of Clinical microbiology, Sahlgrenska University Hospital, Västra Götalandsregionen, Sweden; Simplexia AB, Gothenburg, Sweden; Proteomics Core Facility, Sahlgrenska Academy, University of Gothenburg, Gothenburg, Sweden; Department of Medical Biochemistry and Cell Biology, Institute of Biomedicine, Sahlgrenska Academy, University of Gothenburg, Gothenburg, Sweden

**Author notes:** Corresponding authors: Ebba Könighofer, Rickard Nordén.

**Keywords:** Glycoproteomics, Glycopeptide, Glycoprotein G, Immunogenicity, Vaccine, Neuroinvasion, HSV-2

## Abstract

The function of glycoprotein G (gG-2) of herpes simplex virus 2 (HSV-2) during genital infection is unknown. gG-2 is cleaved into a secreted variant (sgG-2) and a membrane associated variant (mgG-2). This work delineates the glycan profile of mgG-2, and demonstrates that mgG-2 is strongly immunogenic, eliciting glycan dependent humoral and Th1-polarized CD4+ T cell responses in a mouse vaccination model. The N- and O-glycosylation of mgG-2 is important to achieve full protection against HSV-2 infection as immunization with deglycosylated mgG-2 resulted in poor CD4+ T cell activation and viral spread to dorsal root ganglia and spinal cord. Furthermore, an mgG-2 negative HSV-2 mutant virus failed to spread to the dorsal root ganglia and the central nervous system of genitally infected mice, despite viral replication in vaginal cells. Our data demonstrate that mgG-2 is important for HSV-2 propagation and that adaptive immune responses targeting the glycosylated protein can prevent neuronal infection, identifying mgG-2 as a promising vaccine candidate.

## INTRODUCTION

Human herpes simplex virus type 2 (HSV-2) is classified into the *Alphaherpesvirinae* subfamily due to its capacity to establish latency in sensory neurons. HSV-2 infects the genital mucosa and establishes a lifelong latent infection in sensory neurons of dorsal root ganglia (DRG) located in the lumbosacral region of the spine. After reactivation in DRG, the virus is antegradely transported to the periphery, causing genital lesions or asymptomatic shedding of the virus. HSV-2 is also a significant cause of infections within the central nervous system (CNS), arising either during the primary infection or by recurrent reactivation of latent virus (1, 2). In 2020, it was estimated that over 500 million people aged 15–49 years were infected worldwide (3). Two prophylactic vaccine candidates, based on HSV-2 envelope glycoprotein gB-2 and/or gD-2, have reached phase III clinical trials but failed to prevent HSV-2 infection (reviewed by (4)).

HSV-2 encodes 12 envelope glycoproteins responsible for various functions, including viral entry, egress, cell-to-cell spread, and neuronal transport. Like several other envelope glycoproteins, as for example gC, gI and gE (5–8), the glycoprotein G of HSV-2 (gG-2) is dispensable for virus propagation in cell cultures (9). During protein maturation, gG-2 is post-translationally cleaved into a secreted amino-terminal protein (sgG-2) and a carboxy-terminal high mannose-containing intermediate that is further processed by N- and O-linked glycosylation, to constitute the cell-membrane anchored mature gG-2 (mgG-2) (10–15). While the exact glycosylation structures have not been established, the amino-terminal half of mgG-2 is predicted to be a mucin-like region based on the high abundance of the amino acid residues proline, serine, threonine and alanine. Both sgG-2 and mgG-2 stimulate HSV-2 type-specific B- and T-cell responses (16–20), and mgG-2 is widely used as an antigen for type-discriminating serology. Functional studies also show that a sgG-2 peptide functions as a chemoattractant for both monocytes and neutrophils (21), and enhances chemotaxis and chemokine function (22). Furthermore, sgG-2 also modifies nerve growth factor signalling to attract free nerve endings at the site of infection, possibly contributing to novel immune escape mechanisms utilized by HSV-2 (23). While the functional role of mgG-2 in human genital infection remains unclear, data demonstrate its potential as a vaccine candidate in a mouse genital challenge model, suggesting that the immune response against mgG-2 can inactivate important functions of the protein in primary HSV-2 infection (24). An mgG-2 negative HSV-2 strain was shown to spread mostly from cell to cell and had an impaired capacity to release HSV-2 virions into the extracellular medium of the cell culture. In addition, the trapped virions at infected cell membranes were fully virulent and could be released from the cells with sulphated oligosaccharides such as heparin (25). In this work, the role of mgG-2 during *in vivo* infection and the potential use of mgG-2 as a vaccine antigen against HSV-2 infection, was explored. In addition, we probed the role of the N- and O-linked glycans of mgG-2 in shaping the protective immune response in a mouse vaccination model.

To dissect the functional role of mgG-2 *in vivo*, we used an mgG-2 negative mutant and a rescued HSV-2 strain in a genital infection model across two mouse strains. Additionally, we delineated the entire N- and O-linked glycosylation pattern of a recombinant mgG-2 subunit vaccine candidate into discrete glycosylation-profiles and evaluated their respective protective effect against HSV-2 challenge. The results highlight the importance of mgG-2 for viral spread to DRG and to the CNS. As a vaccine antigen, mgG-2 confers protection against a lethal genital challenge with HSV-2 in mice. The protective effect of mgG-2 was attributed to reduced spread of HSV-2 to DRG and CNS in the immunized mice compared to control. Lastly, we showed that glycosylation of mgG-2 is important for its function as a vaccine antigen.

## RESULTS

### Recombinant EXCT4-mgG-2 is decorated with sialylated N- and O-linked glycans

First, we produced a recombinant truncated mgG-2 (EXCT4-mgG-2) in Chinese Hamster Ovary (CHO-K1) cells (Fig. 1A). Using liquid chromatography tandem mass spectrometry (LC-MS/MS) analysis of protease-cleaved glycopeptides, we dissected the glycan profile of the recombinant protein (Fig. 2A-C). Protease cleavage of the EXCT4-mgG-2 prior to LC-MS/MS was performed using alphalytic and pronase. Representative peptides, covering all N-linked consensus sites and 63 out of 70 (90%) possible sites for O-linked glycosylation, were identified and curated for glycan composition and abundance (Fig. S1A and Table S1). Peptide stretches where the fragment spectra evaluation could not confirm the identity of the peptides were excluded from the analysis.

**Figure 1.**
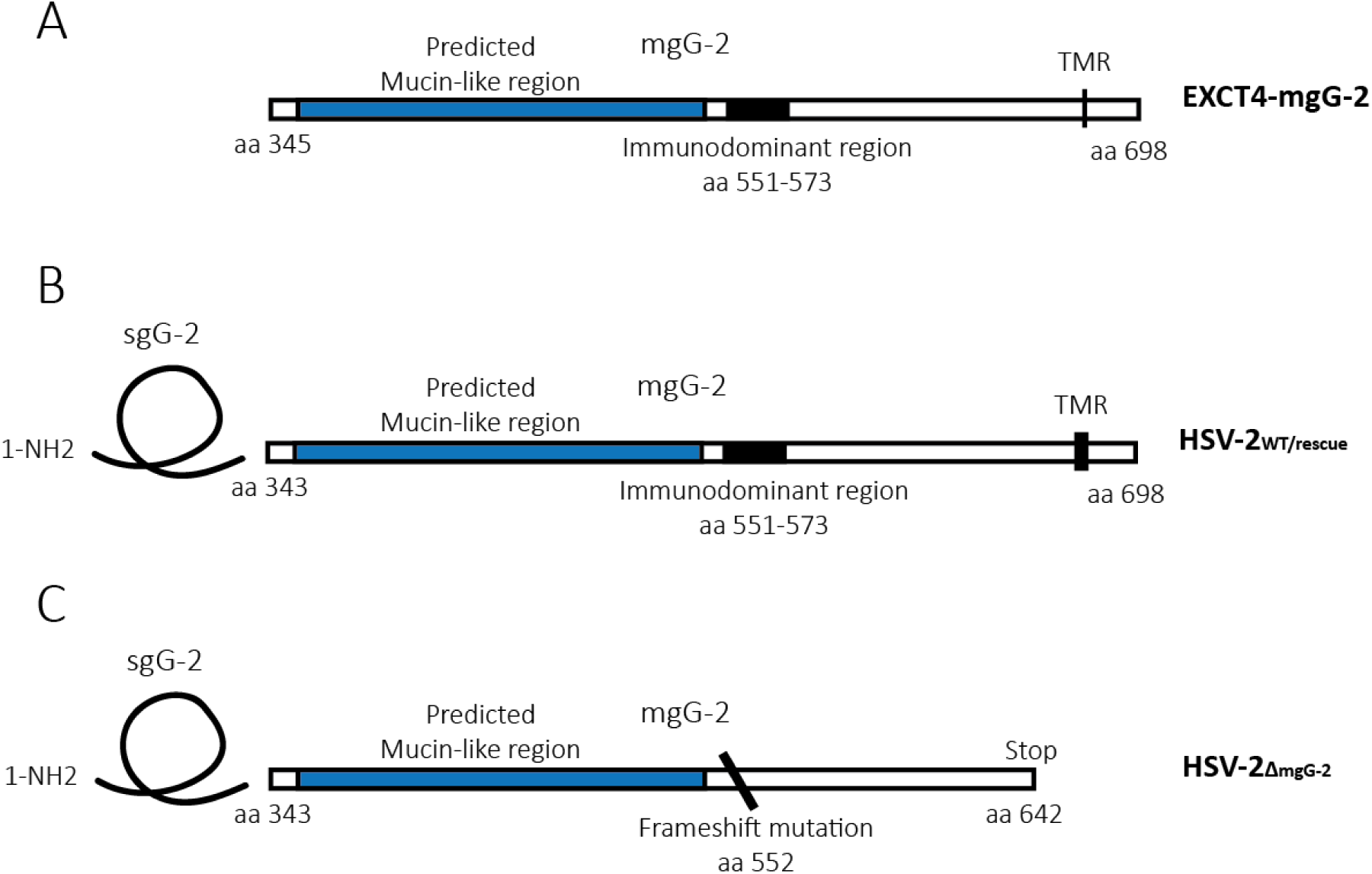
Schematic illustration of the gG-2 protein of HSV-2 and its variants used in the current study. **(A)** The recombinantly produced EXCT4-mgG-2 contains the extracellular region, four amino acids of TMR, and the entire intracellular region. **(B)** The wild type gG-2 protein of HSV-2_WT_ is cleaved generating a secreted portion (sgG-2) and a cell membrane-anchored portion (mgG-2). HSV-2_rescue_ was generated using a marker transfer assay and restores the HSV-2_WT_ genotype. **(C)** The HSV-2_ΔmgG-2_ mutant harbours a frame shift mutation within nucleotides coding for amino acid 552 generating a premature stop at amino acid 642. The immunodominant region for antibodies is marked including the amino acids 551-573. The predicted mucin-like region is indicated. The trans-membrane region is denoted TMR.

**Figure 2.**
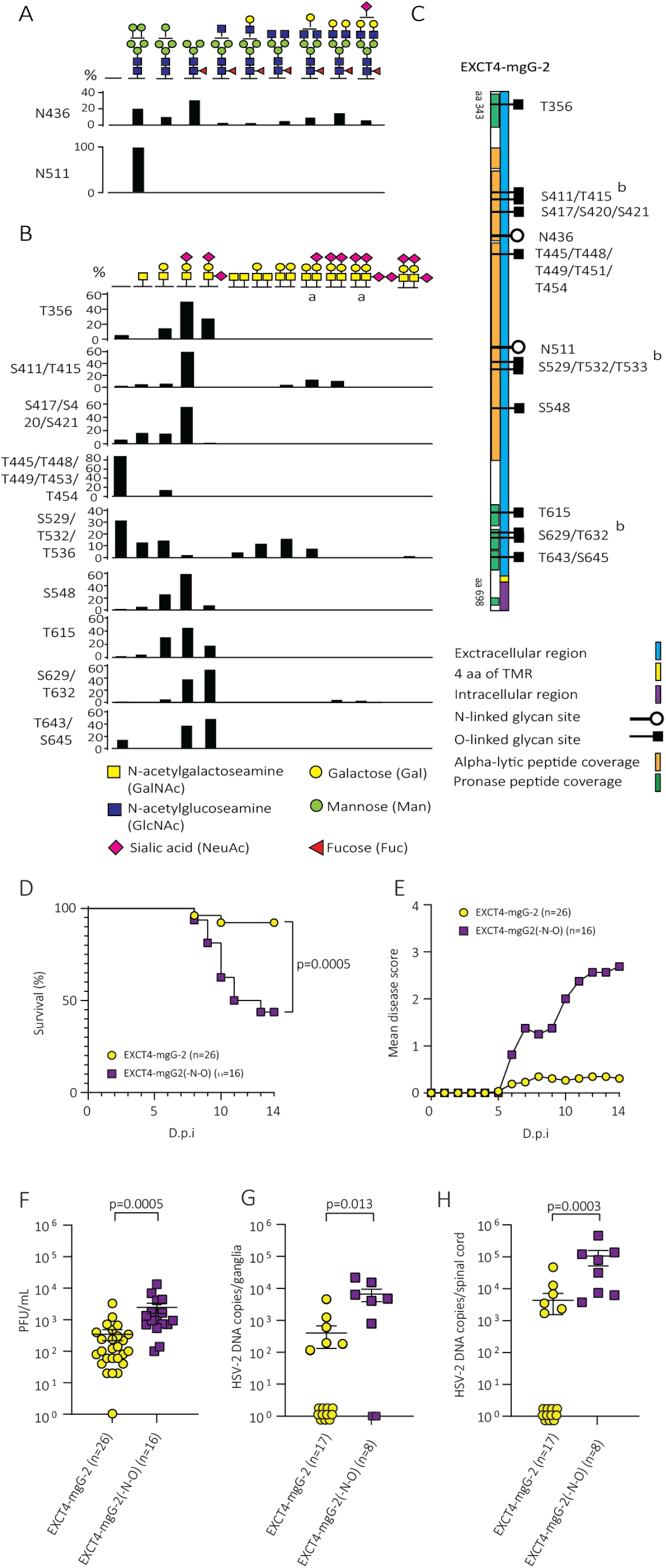
EXCT4-mgG-2 contains multiple N- and O-linked structures which facilitate protection against neuronal spread of HSV-2 in immunized mice. **(A)** Distribution between different glycoforms at each identified glycosite for N-linked glycans. **(B)** Distribution between different glycoforms at each identified glycosite for O-linked glycans. The exact glycosite could not be determined for all O-linked glycans, all possible sites for the respective glycan are stated and indicated with “/”. In the cases where more than one O-linked glycan was observed on the same peptide, two separate glycoforms are present on the same peptide in the schematic drawing. Black bars indicate the percentage of the identified glycoforms on each respective site. Monosaccharide symbols are represented according to the Symbol Nomenclature for Glycans (SNFG) system (54). **(C)** Schematic drawing of EXCT4-mgG-2, including peptide coverage using alphalytic and pronase-treatment, and glycosites for N-linked (black) and O-linked (white) glycosylation. All detected O-linked glycans, regardless of the frequency, are indicated. **(D-E)** C57BL/6 mice were intramuscularly immunized with EXCT4-mgG-2 and EXCT4-mgG-2(−N−O) and genitally challenged with 25 × LD_50_ of HSV-2_WT_. The survival rate and disease score were assessed until 15 d.p.i. **(F)** The number of HSV-2 plaque forming units (PFU) in vaginal washings 2 d.p.i. **(G)** HSV-2 DNA copies per ganglia at 2 d.p.i. **(H)** HSV-2 DNA copies per spinal cord at 2 d.p.i. The detection limit for plaque forming units in vaginal washings was 20 PFU/mL and 40 or 160 HSV-2 DNA copies per 3 ganglia or whole spinal cord respectively. Statistical analysis was performed with the pairwise log-rank (Mantel Cox) test (D), and Kruskal-Wallis test (F-H). D.p.i. - days post infection. Values are expressed as means ± SEM. ^a^ The sialic acid has two possible sites, but the exact position could not be determined. ^b^ More than one O-linked glycan was observed on the same peptide.

EXCT4-mgG-2 contains three sites with consensus sequence Asn-x-Ser/Thr (x = any amino acid, except Pro), which allow for N-linked glycosylation, of these, two (N436 and N511) were identified to be glycosylated (Fig. S1B and Table S2). Site N436 was found to be glycosylated on all identified peptides, primarily by complex type glycans (39.8%) with the most processed structure identified consisted of GlcNAc4Man3Gal2Fuc1NeuAc1, while the oligomannose and paucimannose structures at site N436 were of similar levels (29.8% and 30.4% respectively) (Fig. 2A). Site N511 was found to be exclusively decorated with high mannose (Man5GlcNAc2) (Fig. 2A). Of the 70 possible sites (Ser/Thr) for O-linked glycosylation, 12 were found to carry glycan structures (Fig. 2B). Overall, the most prevalent glycoform was the T-antigen (GalNAc1Gal1), which was found both non-, mono- and di-sialylated. Of note, two of the peptides carrying glycoforms of the N-linked glycan of N436, were also identified as simultaneously carrying an O-linked glycan structure. However, this O-linked structure could not be independently validated when the glycan search was restricted to an O-glycan library alone.

### EXCT4-mgG-2 protection against HSV-2 infection is glycosylation dependent

Next, enzymatic treatment protocols were used to remove specific glycan structures from EXCT4-mgG-2, generating five alternative antigens with 1: N- and O-glycosylated (EXCT4-mgG-2), 2: lacking both N- and O-linked glycans (EXCT4-mgG-2(−N−O)), 3: lacking N-linked glycans (EXCT4-mgG-2(-N)), 4: lacking O-linked glycans (EXCT4-mgG-2(-O)), or 5: lacking sialic acids (EXCT4-mgG-2(-SA)). Glycan trimming was confirmed by western blot and lectin binding-assay (Fig. S2). Next, C57BL/6 mice were immunized intramuscularly three times with respective antigen, followed by genital challenge with 25 × lethal dose 50% (LD_50_) of HSV-2 wild type virus (HSV-2_WT_) and thereafter the survival, disease score, viral replication and spread were monitored. Immunization with EXCT4-mgG-2 conferred protection, showing 93.7% survival 14 days post infection (d.p.i.) (n=26), while immunization with EXCT4-mgG-2(−N−O) resulted in a survival of 43.8% (p=0.0025, n=16) (Fig. 2D). The mean disease score for mice immunized with EXCT4-mgG-2 was 0.3 at 14 d.p.i., as compared to a mean disease score of 2.7 for the mice immunized with EXCT4-mgG-2(−N−O) (Fig. 2E). The viral load was higher in vaginal washes from mice that were immunized with the deglycosylated antigen EXCT4-mgG-2(−N−O) compared to mice that received EXCT4-mgG-2 (p=0.0005) (Fig. 2F). Mice immunized with EXCT4-mgG-2(−N−O) presented with higher levels of HSV-2 DNA in DRG than mice immunized with EXCT4-mgG-2, 48 hours post infection (h.p.i.) (p=0.013) (Fig. 2G). Also, the HSV-2 DNA copy numbers in the spinal cord were significantly higher in mice immunized with EXCT4-mgG-2(−N−O) (p=0.0003) (Fig. 2H), indicating poorer protection against viral spread in neuronal tissue. In contrast, immunization with EXCT4-mgG2(-N) (n=8), EXCT4-mgG-2(-O) (n=8) or EXCT4-mgG-2(-SA) (n=8), all conferred protection against HSV-2_WT_ and the mice showed similar disease score as mice immunized with EXCT4-mgG-2 (Fig. S3A-B). In addition, no significant impairment in the protective effect on viral spread in neuronal tissue was observed compared to EXCT4-mgG-2 (Fig. S3C-D). In conclusion, negative effect on the immune response occurs when both the N- and O-linked glycans are removed from the antigen.

### EXCT4-mgG-2 generates a glycosylation dependent adaptive immune response

To investigate the T-cell responses, spleens were harvested 10 days after the final immunization of C57BL/6 mice, and the splenocytes were stimulated with a peptide pool consisting of overlapping 15-mers spanning the entire EXCT4-mgG-2 protein sequence. The cells were then subjected to fluorospot analysis, which showed that interferon gamma (IFN-γ), interleukin 2 (IL-2) and tumour necrosis factor alpha (TNF-α) were significantly more expressed in splenocytes from EXCT4-mgG-2-immunized mice compared to those from EXCT4-mgG-2(−N−O)-immunized mice (Fig. 3A). Moreover, the frequency of splenocytes expressing all three cytokines simultaneously was higher in the EXCT4-mgG-2 group (Fig. 3B). Splenocytes from mice immunized with EXCT4-mgG-2(−N−O) also showed elevated levels of all three cytokines compared to the mock immunized mice (Fig. 3A-B). Next, splenocytes were stimulated with the intact glycosylated EXCT4-mgG-2 antigen and subjected to fluorospot analysis. Splenocytes from both immunized groups, EXCT4-mgG-2 and EXCT4-mgG-2(−N−O), showed higher expression of both INF-γ and IL-2 compared to the mock controls (Fig. 3C). Notably, only IL-2 expression was significantly higher in EXCT4-mgG-2 group compared to the EXCT4-mgG-2(−N−O) group. Then, splenocytes were stimulated with the 15-mer peptide pool, and intracellular INF-γ expression in CD4+ and CD8+ T cells was assessed by flow cytometry (Fig. S4). A fraction of CD8+ T cells from mice immunized with EXCT4-mgG-2 and EXCT4-mgG-2(−N−O) showed intracellular staining for INF-γ, compared to CD8+ T cells from the mock immunized mice (Fig. 3D). Only CD4+ T cells from mice immunized with the EXCT4-mgG-2 antigen demonstrated intracellular INF-γ expression (Fig. 3D). When combining both CD4+ and CD8+ T cells responses, only the EXCT4-mgG-2 immunized group exhibited a clearly increased number of INF-γ reactive cells (Fig. 3D). We next determined if the CD4+ T cell response induces a glycosylation dependent antibody response against EXCT4-mgG-2. Serum samples were collected 14 days following the third intramuscular immunization with EXCT4-mgG-2 or EXCT4-mgG-2(−N−O) and the IgG1 and IgG2c levels were determined. Immunization with EXCT4-mgG-2 generated lower titres of IgG1 compared to EXCT4-mgG-2(−N−O) while there was no difference in IgG2c titres (Fig. 3E) corroborating that glycosylation is important for mounting a Th1 response. Furthermore, the serum samples were investigated for reactivity towards EXCT4-mgG-2 and EXCT4-mgG-2(−N−O) in an ELISA. Antibodies in serum from EXCT4-mgG-2(−N−O) immunized mice showed reduced reactivity towards EXCT4-mgG-2, while antibodies in serum from EXCT4-mgG-2 immunized mice equally well recognized both EXCT4-mgG-2 and EXCT4-mgG-2(−N−O) (Fig. 3F). To compensate for the antibody reactivity against the glycosidases used in the deglycosylation of EXCT4-mgG-2(−N−O), mice were immunized with an injection fluid containing the glycosidases but lacking EXCT4-mgG-2. Sera from these mice were analysed by ELISA for reactivity against EXCT4-mgG-2 and EXCT4-mgG-2(−N−O) (Fig. S5). The mean antibody titres obtained from these control animals were subtracted from the titres of mice immunized with EXCT4-mgG-2(−N−O), depicted in Figure 3F. Taken together, it appears that CD4+ T cells can differentiate between epitopes within mgG-2 depending on the glycosylation status, indicating glycosylation dependent recognition by CD4 + T cells that can inactivate important functions of the HSV-2 associated mgG-2 upon subsequent infection, thereby preventing neuronal spread.

**Figure 3.**
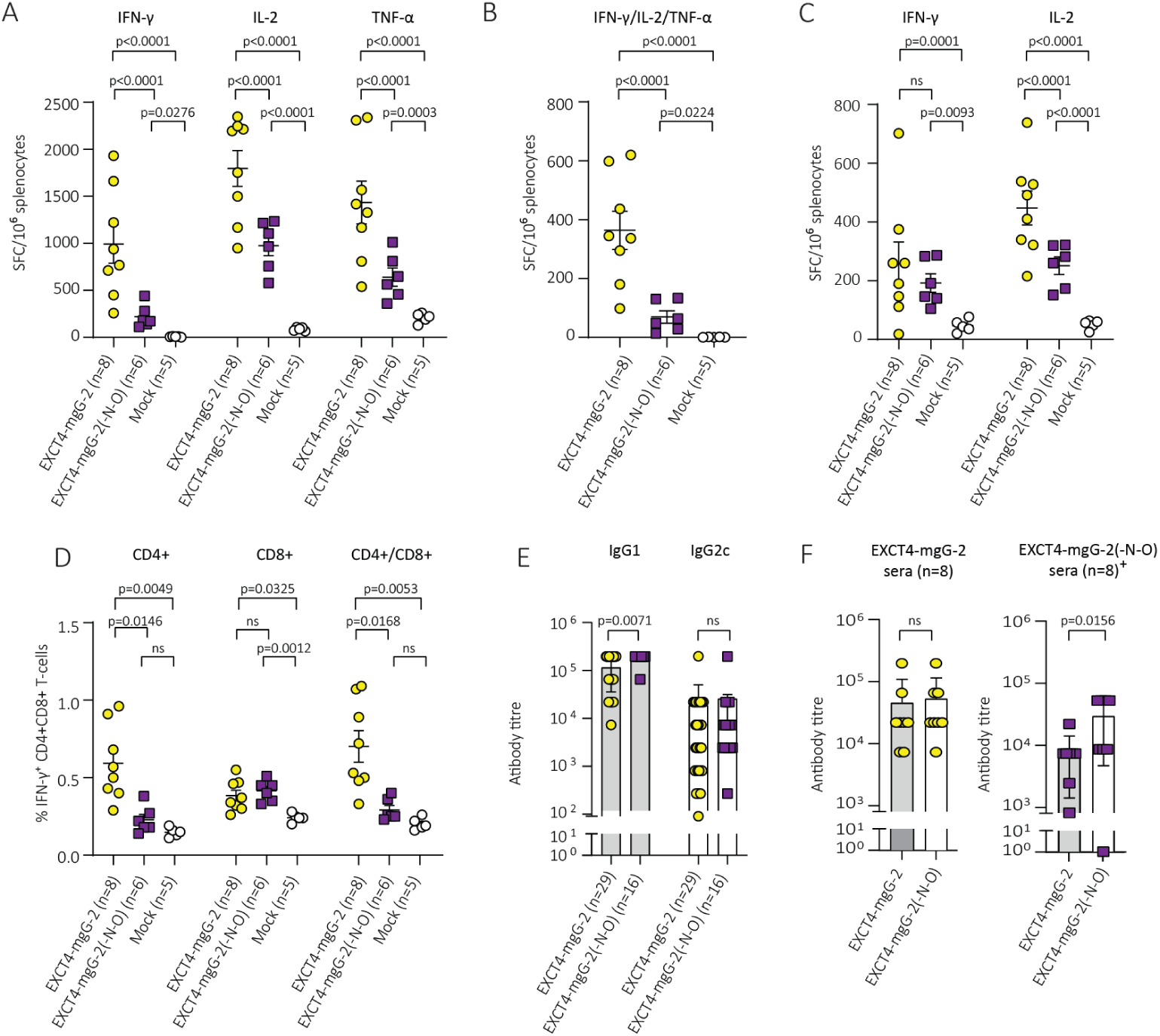
EXCT4-mgG-2 induces Th1 polarization after immunization, a response which is attenuated in the absence of glycosylation. **(A)** Flourospot analysis of splenocytes harvested after immunization and stimulated with a pool of overlapping peptides (15-mers) covering the entire EXCT4-mgG-2 sequence, showing INF-γ, IL-2 and TNF-α production. **(B)** Fluorospot analysis showing the combined expression of INF-γ, IL-2 and TNF-α in splenocytes stimulated with the 15-mer peptide pool **(C)** Fluorospot analysis showing INF-γ and IL-2 expression in splenocytes stimulated with the N- and O-glycosylated EXCT4-mgG-2 recombinant protein **(D)** Percentage of INF-γ positive CD4+ and CD8+ T cells from splenocytes stimulated with the 15-mer peptide pool. The splenocytes were harvested 14 days after the third immunization with EXCT4-mgG-2, EXCT4-mgG-2(−N−O) or mock and stimulated with the peptide pool overnight. **(E)** IgG1 and IgG2c levels in serum samples collected 14 days after the third intra muscularly (i. m.) immunization with EXCT4-mgG-2 or EXCT4-mgG-2(−N−O). **(F)** Serum antibody reactivity against EXCT4-mgG-2 and EXCT4-mgG-2(−N−O) was assessed in serum samples collected 14 days following immunization with EXCT4-mgG-2 or EXCT4-mgG-2(−N−O). ^+^ For immunization group EXCT4-mgG-2(−N−O), the antibody reactivity was corrected by subtracting values corresponding to glycosidase-specific binding in serum from mice immunized with the glycosidase enzymes in absence of EXCT4-mgG-2. SFC = Spot forming cells. Statistical analysis was performed with Saphiro-wilk test and Tukeýs multiple comparison test (A-D), Kruskal-Wallis test (E) or Mann-Whitney test (F). Values are expressed as means ± SEM.

### The HSV-2_ΔmgG-2_ mutant produces sgG-2

To further explore the function of mgG-2 in facilitating neuronal spread we utilized HSV-2 mgG-2 negative (HSV-2_ΔmgG-2_), HSV-2 wild type (HSV-2_WT_) and HSV-2 rescued (HSV-2_rescue_) viral strains (Fig. 1B and C). First, the production of sgG-2 for HSV-2_ΔmgG-2_, HSV-2_rescue_, and HSV-2_WT_ was determined. African green monkey kidney cells (GMK-AH1) were infected at a multiplicity of infection (MOI) 1 of respective viral strain, and the supernatants were collected when complete cytopathic effect was observed. The supernatants were normalized to the same amount of produced virus (PFU/mL). Western blot analysis indicated as expected that sgG-2 (44 kDa) are produced for all three HSV-2 strains (Fig. S6). The lack of expression of mgG-2 in the HSV-2_ΔmgG-2_ viral strain was previously verified (25).

### HSV-2_ΔmgG-2_ infects vaginal tissue but presents low mortality and disease scores

C57BL/6 mice were infected intravaginally with 10 × LD_50_ of HSV-2_WT_, HSV-2_ΔmgG2_ or HSV-2_rescue_. Survival (Fig. 4A) and disease score (Fig. 4B) were observed for 21 d.p.i. In mice infected with HSV-2_WT_ or HSV-2_rescue_, symptoms appeared 6 d.p.i. and presented a progressive genital and systemic infection and were all euthanized. In contrast, all mice infected with HSV-2_ΔmgG-2_ survived (Fig. 4A) and presented no or limited inflammation in vagina, vulva and perineal region reflected by low disease scores (Fig. 4B). No clinical signs of spread to the autonomic neurons (constipation and urine retention) were observed in the mice infected with HSV-2_ΔmgG-2_. Next, the more HSV-2 susceptible DBA/2 mice were infected with 125 × LD_50_. The outcome of the infection with HSV-2_WT_ and HSV-2_rescue_ was similar, and all mice were euthanized (Fig. 4C and D). After infection with HSV-2_ΔmgG-2_, 25 of 29 (86 %) mice survived. Although four mice succumbed, the infection was more prolonged, and the mice were euthanized between 12 and 16 d.p.i. (Fig. 4C and D). To exclude that the HSV-2_ΔmgG-2_ had reverted to the HSV-2_WT_ phenotype, spinal cord DNA from one euthanized DBA/2 mouse was prepared for HSV-2 gG-2 gene sequencing as described (26). The sequence presented the identical frameshift mutation in the mgG-2 gene as described for HSV-2_ΔmgG-2_. In conclusion, HSV-2_ΔmgG-2_ shows greatly reduced capacity to promote genital and neurological disease.

**Figure 4.**
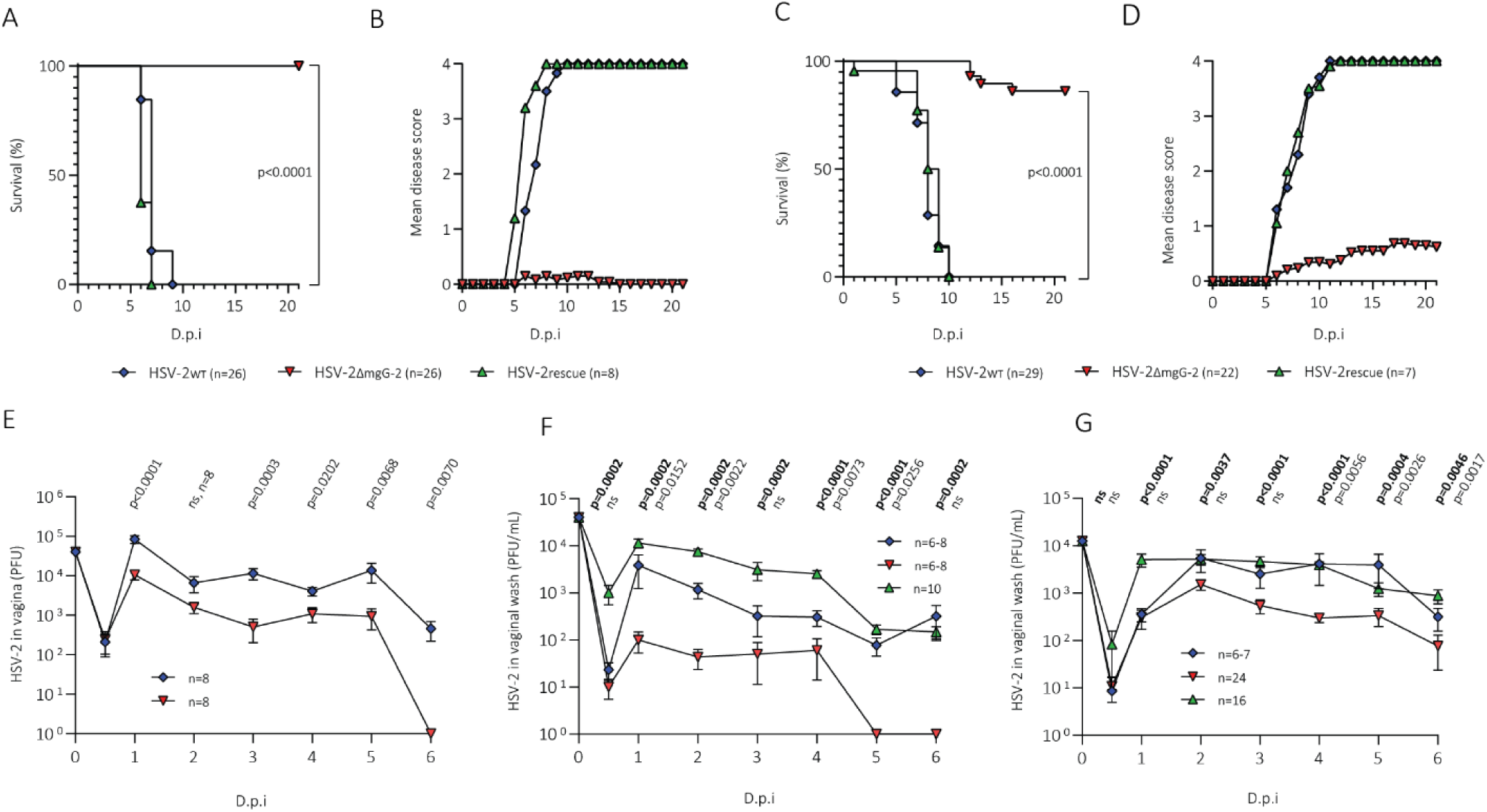
HSV-2_ΔmgG-2_ infection induces low genital disease scores and low mortality. In addition, HSV-2_ΔmgG-2_ infects and replicate in the genital tract but show impaired release from the surface of vaginal epithelial cells. **(A-B)** C57BL/6 mice were infected intravaginally with 10 × LD_50_ of HSV-2_WT_, HSV-2_ΔmgG-2_ or HSV-2_rescue_. The survival rate and disease score were followed until day 21. **(C-D)** DBA/2 mice were infected intravaginally with 125 × LD_50_ of HSV-2_WT_, HSV-2_ΔmgG-2_ or HSV-2_rescue_. **(E)** Vaginas from C57BL/6 mice were excised at different timepoints after genital infection with 10 × LD_50_ of HSV-2_WT_ or HSV-2_ΔmgG-2_ and HSV-2 plaque forming units (PFU) were assessed. The total PFU production between day 1 and day 2 was calculated by area under curve method (AUC). **(F)** The PFU content in vaginal washings of C57BL/6 infected mice. **(G)** The PFU content in vaginal washings from DBA/2 mice infected with 125 × LD_50_ of HSV-2_WT_, HSV-2_ΔmgG-2_ or HSV-2_rescue_. PFUs (detection limit 40 PFU/mL) are expressed as means ± SEM. Statistical analysis is performed with pairwise Log-rank (Mantel-Cox) test (A and C) or Mann-Whitney test (E-G). P-values are calculated for HSV-2_rescue_ versus HSV-2_ΔmgG-2_ indicated in bold and for HSV-2_WT_ versus HSV-2_ΔmgG-2_ indicated in normal script (F and G). Data are calculated from two to three independent experiments. D.p.i. - days post infection.

Next, C57BL/6 mice were infected with 10 × LD_50_ of HSV-2_WT_ or HSV-2_ΔmgG-2_ and the vagina was excised 12 h.p.i. or 1, 2, 3, 4, 5 or 6 d.p.i. For both HSV-2_WT_ and HSV-2_ΔmgG-2_ the number of infectious particles declined approximately 100 times, compared to the input levels, at 12 hours past infection (h.p.i.). (Fig. 4E). The peak in viral load was reached 24 h.p.i. for both strains. At 6 d.p.i. with HSV-2_ΔmgG-2_ no virus was detectable and HSV-2_ΔmgG-2_ presented lower viral loads at all time points from day 1 and onwards.

The HSV-2 infection was estimated in vaginal washes at the same time-points after infection also including the HSV-2_rescue_ strain. Although at lower levels, similar kinetics was described as for the vaginal infection. The input virus was readily cleared within 12 h.p.i. and the peak in viral load was obtained 1 d.p.i. (Fig. 4F). The differences between HSV-2_rescue_ and HSV-2_ΔmgG-2_ were statistically significant for all time points after infection. In cultured primary vaginal epithelial cells derived from C57BL/6 mice, HSV-2 was shown to be shed mostly from the apical surface (27). The viral load in the vaginal washes can therefore be used as marker of release of extracellular HSV-2 in the vaginal infection before virus infected epithelial cells are sloughed off. Based on the area under curve method for estimation of the viral load between day 1 and day 2, HSV-2_WT_ produced 7.6 times more PFU as compared with HSV-2_ΔmgG-2_ in vaginal tissue while in vaginal washes the difference was 77 times (p=0.01), (Fig. 4E and F).

The more sensitive DBA/2 mice were infected with 125 × LD_50_, and infectious particles were measured in the vaginal washes (Fig. 4G). The sum of mean values of infectious particles for 12 h.p.i. and 1-6 days past infection (d.p.i.) in the vaginal washes was 29,223 PFU for HSV-2_WT_, 39,626 PFU for HSV-2_rescue_, and 5,945 PFU for HSV-2_ΔmgG-2_. The differences between HSV-2_rescue_ and HSV-2_ΔmgG-2_ were statistically significant for days 1-6. As the production of infectious viral particles was measured only in vaginal washes no ratio between PFU in vagina versus vaginal washes could be calculated.

### Reduced spread of HSV-2_ΔmgG-2_ in serum, genital lymph nodes and neuronal tissue

After infection of C57BL/6 mice with 10 × LD_50_ of HSV-2_WT_ the viral DNA genome copy numbers in serum reached a plateau 1 d.p.i., which lasted until 4 d.p.i. whereafter the viral load increased almost 10 times (Fig. 5A). In contrast, after infection with 10 × LD_50_ of HSV-2_ΔmgG-2_ viral DNA increased more slowly and reached lower levels as compared with the HSV-2_WT_ and was cleared 6 d.p.i. Viral load was also measured after infection with HSV-2_rescue_, which presented similar values as for HSV-2_WT_ 6 d.p.i. (Fig. 5A). These data show, also including results from the vaginal infection, that there is a clear difference in the kinetics of the infection between HSV-2_WT_ and HSV-2_ΔmgG-2_. Thus, in the initial stage up to day 4-5, the difference in the viral load is moderate, while after this period the infection with HSV-2_ΔmgG-2_ is rapidly cleared. In DBA/2 mice, HSV-2 DNA was measured in serum 6 d.p.i. Similar levels were detected for HSV-2_WT_ and for HSV-2_rescue_ as described for the C57BL/6 mice. One DBA/2 mouse of eight, infected with HSV-2_ΔmgG-2_, presented a value of 747 viral DNA copy numbers per mL serum (Fig. 5B).

**Figure 5.**
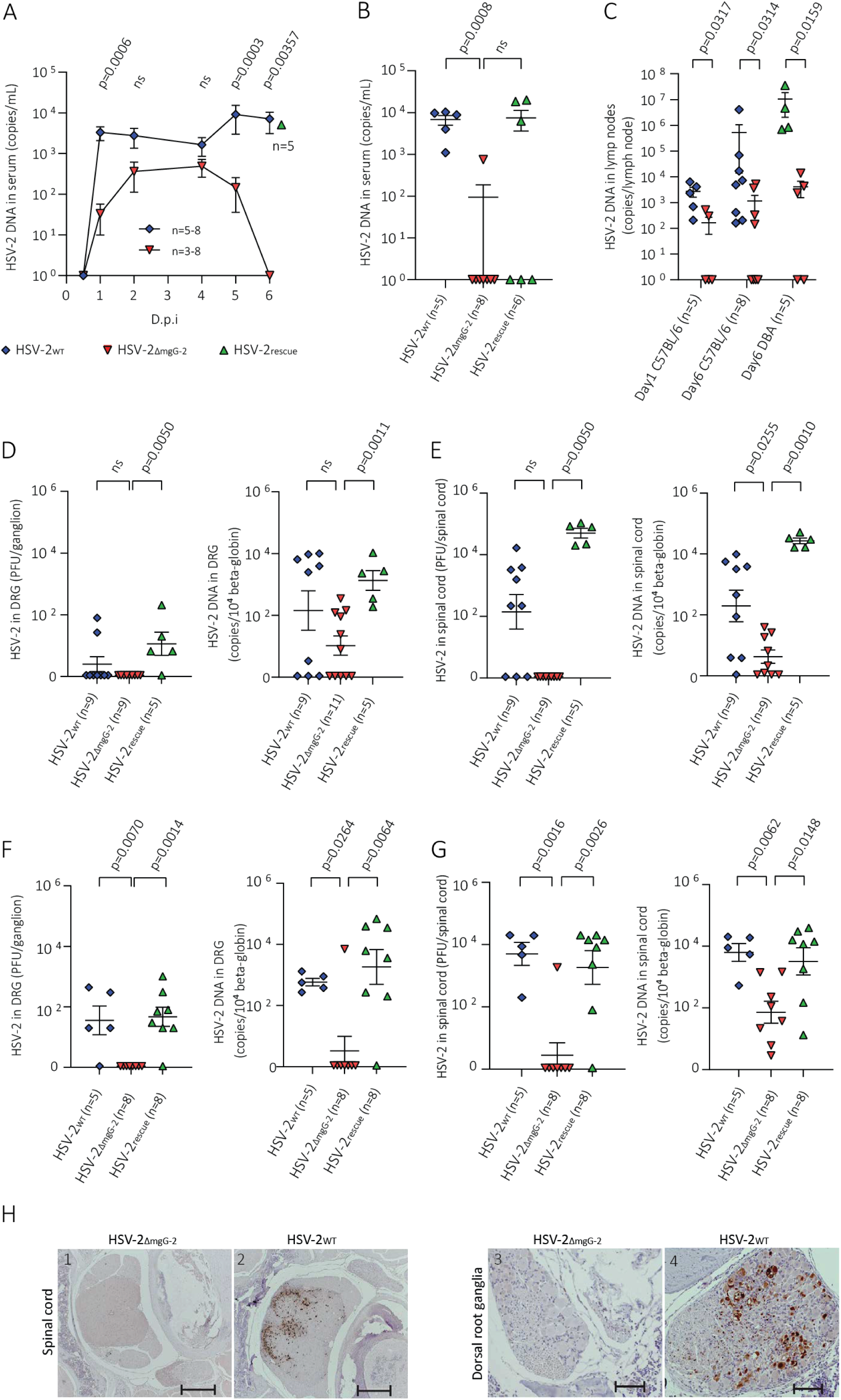
Assessment of the spread of HSV-2 to blood, genital lymph nodes, dorsal root ganglia and spinal cord, showing impaired capacity for neuronal spread of HSV-2_ΔmgG-2_. **(A)** C57BL/6 mice were infected with 10 × LD_50_ of HSV-2_WT_ or HSV-2_ΔmgG-2_, and HSV-2 DNA was calculated at 1-6 d.p.i. in serum. Sera from 5 C57BL/6 mice infected with the HSV-2_rescue_ were analysed at day 6 (marked in green) **(B)** DBA/2 mice were infected with 125 × LD_50_ of HSV-2_WT_, HSV-2_ΔmgG-2_ or HSV-2_rescue_, and HSV-2 DNA content in serum was calculated at 6 d.p.i. **(C)** C57BL/6 mice were infected with 10 × LD_50_ of HSV-2_WT_ or HSV-2_ΔmgG-2_ and HSV-2 DNA was calculated in genital lymph nodes at 1 or 6 d.p.i. **(D-E)** C57BL/6 mice were infected with 10 xLD_50_ and **(F-G)** DBA/2 mice were infected with 125 × LD_50_ of HSV-2_WT_, HSV-2_ΔmgG-2_ or HSV-2_rescue_. Infectious HSV-2 and HSV-2 DNA copy numbers were analysed 6 d.p.i. (D and F) per single dorsal root ganglion and (E and G) in the whole spinal cord. **(H)** Paraffin sections from the vertebral column of DBA/2 mice 6 d.p.i. infection with (panel 1 and 3) HSV-2_ΔmgG-2_ or (panel 2 and 4) with HSV-2_rescue_. HSV-2 antigens (brown) were visualized using a polyclonal rabbit anti-HSV-2 serum. Panel 1-2 show the spinal cord at the thoracic level and panel 3-4 show the dorsal root ganglia at the lumbosacral level from the same animal. After infection with HSV-2_rescue_ several neurons and glial cells in the dorsal horns and neurons and satellite cells in the ganglia are infected. Bars: 1-2; 500 μm, 3-4; 100 μm. HSV-2 DNA copies are expressed per mL serum (detection limit 100 copies) or per lymph node (detection limit 80 copies). Values are expressed as means ± SEM. HSV-2 DNA copy numbers were standardized to 10^4^ beta-globin genes. The virus detection limit was 40 PFU or 160 HSV-2 DNA copies per 3 ganglia or whole spinal cord. Statistical analysis is performed with Mann-Whitney test. Data are calculated from one to two independent experiments for each mouse strain except for the immunohistochemistry data where one experiment representative of the two performed is shown.

In C57BL/6 infected mice, viral DNA was detected in genital lymph nodes 1 d.p.i. with an increase of the viral load for HSV-2_WT_ to 4 × 10^5^ viral DNA copy numbers 6 d.p.i., as compared with 1 × 10^3^ copy numbers after infection with HSV-2_ΔmgG-2_. In DBA/2 infected mice, the mean viral load per lymph node 6 d.p.i. for HSV-2_rescue_ was 1 × 10^7^ HSV-2 DNA copy numbers as compared with 5 × 10^3^ after infection with HSV-2_ΔmgG-2_ (Fig. 5C).

Next, C57BL/6 mice were infected with 10 × LD_50_ and DBA/2 mice with 125 × LD_50_ of HSV-2_WT_, HSV-2_ΔmgG-2_ and HSV-2_rescue_. DRG and spinal cord were removed 6 d.p.i., and infectious virus was assayed by a plaque assay and HSV-2 DNA was quantified by real-time PCR. For most animals infected with HSV-2_WT_ or HSV-2_rescue_, viral DNA was detectable in both the DRG and the spinal cord 6 d.p.i. (Fig. 5D and E). These data are similar as those described for progression of the infection in mice after intravaginal infection (28). The most important finding which explains the high survival rate of mice after infection with HSV-2_ΔmgG-2_, was that the viral load in DRG and spinal cord was significantly reduced. After infection of C57BL/6 mice with HSV-2_ΔmgG-2_ no infectious viral particles were detected in DRG and spinal cord. After infection of DBA/2 mice with HSV-2_ΔmgG-2_, infectious virus was only detected in the spinal cord in one mouse and significantly lower values HSV-2 DNA were detected in both DRG and spinal cord as compared with infection with HSV-2_WT_ or HSV-2_rescue_ strains (Fig. 5F and G). The lack of infectious HSV-2 and production of viral proteins in DRG and spinal cord after infection with HSV-2_ΔmgG-2_ were also confirmed with an immunohistochemistry technique using a rabbit polyclonal anti-HSV-2 serum for detection (Fig. 5H).

## DISCUSSION

Although the functions of several of the envelope glycoproteins of HSV-2 have been defined, the role of mgG-2 during the *in vivo* infectious cycle has still to be clarified. Using a genital infection model in two mice strains, we show that HSV-2_ΔmgG-2_ replicates in the vagina but the spread of virus from vagina to blood, to genital lymph nodes, to sensory ganglia, to spinal cord, and to genital autonomic neurons, was severely impaired in the absence of mgG-2. After infection with HSV-2_ΔmgG-2_ infectious viral particles in the vaginal lumen were initially reduced 10 times as compared with HSV-2_WT._ These observations agree with impaired release of infectious HSV-2_ΔmgG-2_ particles in mouse fibroblast L-cell cultures where extracellular viral particles were approximately 10 times lower as compared with HSV-2_WT_ (29). However, when the infective dose of HSV-2_ΔmgG-2_ was increased to 125 × LD_50_ PFU in DBA/2 mice the content of infectious HSV-2 in the vaginal washes was similar at day 2 as for HSV-2_WT_ using the infective dose of 10 × LD_50_ PFU in C57BL/6 mice. Despite this increase, only 4 of 29 DBA/2 mice died while all mice died infected with HSV-2_WT_, suggesting that impaired spread of virus from infected vaginal epithelia to sensory neurons is most important.

In the initial infection HSV uses gB, gC and gD to attach to cellularly produced glycosaminoglycans (GAGs) on the cell membrane. After attachment, the gD binds to a cell receptor for entry. In murine keratinocytes nectin-1 was shown to be the primary receptor (30). Using knock-out C57BL/6 mice lacking the cellular receptor herpes virus entry mediator (HVEM) or nectin-1 or both, Tayler et al., showed that HVEM or nectin-1 was required for infection of the vaginal epithelium after genital infection but neither of the receptors were essential for spread of HSV-2 to DRG and spinal cord (31). After entry and replication, progeny virus are released by exocytosis as individual virions (32, 33) . The last phase of the infection is the release of infectious virions from infected cells for further spread. To avoid that GAGs block the release of virions by binding to the HSV attachment proteins, HSV has developed counteractive strategies. For example, Hadigal et al., showed in HSV-1 infected cells that the levels of cell membrane bound heparan sulfate (HS) GAGs were substantially decreased by up-regulation of the host enzyme heparanase-1, cleaving the HS on the cell surface (34).

In an earlier report, HSV-2_ΔmgG-2_ was shown to be impaired in egress and that mature infectious virus was accumulated on cell infected membranes in different cell lines but could effectively be released using GAG-mimicking oligosaccharides. Using total internal reflection fluorescens microscopy and a single particle tracking analysis of virions to surface bound chondroitin sulfate (CS) GAGs, HSV-2_ΔmgG-2_ virions presented extensive binding to CS which resulted in limited mobility of the virions on the CS surfaces as compared to HSV-2_WT_. A conclusion was that mgG-2 balances the interaction between the virion and GAGs present at virus infected cell membranes to enable egress (29). In the present study HSV-2_ΔmgG-2_ show impaired egress from vaginal epithelial cells which could contribute to the high survival rate in two mouse strains after genital infection with HSV-2_ΔmgG-2_. However, we cannot exclude that mgG-2 has additional functions in the neuronal infection, for example facilitating entry into the nerve endings. As no cellular receptor binding to mgG-2 and no structure of the protein has been defined, further studies are warranted to discover cellular or viral binding partners to mgG-2 which would aid further development of a vaccine.

Another aspect with relevance for vaccine development is the impact of post translational glycosylation of viral surface proteins. To address the knowledge gap regarding the impact of glycosylation of the putative mucin like domain of mgG-2, we defined the entire glycosylation profile of a recombinantly expressed mgG-2 and constructed several glycan modulated vaccine candidates. We showed that immunization with EXCT4-mgG-2 elicited a robust adaptive immune response including Th1-polarized cells that produced IL-2 and INF-γ, as compared to immunization with EXCT4-mgG-2(−N−O), suggesting enhanced T helper cell function that can contribute to B cell help and class switching in favour of IgG2c. Thus, glycosylation of EXCT4-mgG-2 may bias antigen presentation towards MHCII peptides that favour Th1 polarization. This is interesting since glycans can, either on their own or in combination with the protein backbone, constitute B- and T- cell epitopes and thereby enhance antigenicity (35). Previously, a 10 amino acid peptide within mgG-2, carrying a O-linked glycan at threonine 504 (T504), was reported to be immunodominant in a glycosylation dependent manner and recognized by HSV-2 seropositive patient serum samples only in the presence of the glycan structure (36). We could not identify O-linked glycosylation at T504 in EXCT4-mgG-2 but showed that there was a glycosylation dependent reactivity of serum from immunized mice. This suggests that the glycans may contribute directly to glycoepitope formation, however, the fact that spleenocytes from EXCT4-mgG-2-immunized mice showed a stronger response to the unglycosylated peptide pool makes it difficult to reconcile with the presence of T-cell specific glycoepitopes. The dampened response towards the whole glycosylated antigen might be explained by delayed display by the antigen-presenting cells (APCs) as the whole antigen must be processed into shorter peptides before it can be integrated into the MHC-peptide complex. An alternative explanation to the superior efficiency of EXCT4-mgG-2 could be that the glycans focus the response towards immunodominant epitopes in unglycosylated peptide stretches of the antigen, thus, that glycans alter the immune focus by shielding competing epitopes as was recently suggested for HSV-1 glycoprotein B (37). Studies of glycosylation as antigen-dependent factors contributing to the selection of T cell epitopes is scarce but it has been described for HIV-1 glycoprotein gp120 and it was further suggested that epitopes selection was enhanced in protein regions characterised by high accessibility and low stability (38, 39). A third possibility could be that the glycans affect antigen uptake by interaction with glycan-binding C-type lectin receptors on APCs and thereby facilitating a more efficient antigen presentation (40). These possible explanations are not mutually exclusive and thus could act synergistically to generate the enhanced IL-2 and INF-γ response observed for EXCT4-mgG-2 compared to EXCT4-mgG-2(−N−O).

Furthermore, Görander et al. have, using INF-γ-gene knockout mice (KO) on a C57BL/6 background, shown that INF-γ produced by CD4+ T cells are essential to generate protection after immunization with glycosylated native mgG-2 with CpG as adjuvant followed by genital challenge (24). In addition, B cell KO mice, immunized with glycosylated mgG-2 and CpG and alum as adjuvants, presented lower survival rate and higher vaginal viral titers, as compared with vaccinated B-cell KO mice which received passive transfer of immune serum from vaccinated C57BL/6 mice (41). Interestingly, in both studies anti-mgG-2 antibodies presented no neutralizing capacity while all sera exerted antibody-dependent cellular cytotoxicity (ADCC) and complement-mediated cytolysis activity. Here we show that EXCT4-mgG-2 induce protection against genital and neurological disease as well as mortality. The favorable outcome can be explained by low or absent levels of infectious viral particles in spinal cord suggesting that the immune responses against EXCT4-mgG-2 can inhibit the spread of HSV-2 from the vaginal epithelial cells into the sensory neurons, or alternatively clear the infection in already infected neurons by INF-γ producing CD4+ T cells as described for HSV-1 infection in a murine model (42).

Our detailed characterization of the glycan content of EXCT4-mgG-2, contradict the notion that mgG-2 holds densely clustered O-linked glycans in its predicted mucin-like region (Fig. 1). When the protein is recombinantly expressed in CHO-cells, 12 dispersed sites harbour O-linked glycans, mainly of sialylated Tn type, in addition to the expected N-linked glycans. In agreement with this, it was previously reported that gG-2 from HSV-2 strain 333 harboured 18 O-linked glycans when the virus was propagated in African green monkey cells (Vero), of which 4 were situated in sgG-2 and 14 dispersed throughout mgG-2, while the N-linked glycans were not evaluated (43). When examining the occupied glycosites within EXCT4-mgG-2 we found 6 O-glycans that showed a complete overlap with what Iversen et al. observed for mgG-2 from HSV-2 grown in Vero cells (43). Thus, it appears that the number of O-linked glycans is similar but there is only partial overlap when considering the individual glycosites. The discrepancy in glycan occupancy can possibly be attributed to cell specific differences in expression of glycosyltransferases e.g. CHO-cells lack α2-6 sialyltransferase expression, but more importantly exhibit limited expression of a subset of GalNAc-transferases, that could negatively affect site occupancy in addition to alterations in glycan structure heterogeneity (44, 45). For example, the glycan occupancy of glycoprotein E from varicella zoster virus (VZV) was less dense when the protein was expressed in CHO cells, compared to when expressed in primary human fibroblasts (46, 47). Here we show that glycosylation of mgG-2 contributes to generating a more efficient humoral immune response upon immunization; the mice that received the EXCT4-mgG-2 antigen presented a significantly higher survival and a lower disease score compared with the mice that received the deglycosylated antigen EXCT4-mgG-2(-N-O). With the amino acid backbone of the recombinant antigens being identical, we attribute the reduced survival of the EXCT4-mgG-2(−N−O) immunized mice to the absence of glycosylation. In contrast, removal of N-linked, O-linked, or sialic acid glycan structures alone, presented no effect on either mortality or morbidity.

A limitation of this study is that we did not identify individual T cell epitopes and could therefore not assess the impact of individual glycan structures on the above-mentioned effects on the adaptive immune response. Also, in this work, deglycosylated variants of EXCT4-mgG-2 were obtained by enzymatic treatment, and it is expected that residual glycan structures are present on a fraction of the proteins. Consequently, it cannot be ruled out that the partial protective effect observed in mice immunized with EXCT4-mgG-2(−N−O) comes from incompletely deglycosylated EXCT4-mgG-2. Thus, the survival rate of mice immunized with EXCT4-mgG-2(−N−O) antigen may be overestimated. Attempts were made to produce recombinantly expressed mgG-2 in an *in vitro* translation *E. coli* system that would generate a protein devoid of glycan structures, but, despite great efforts, isolation of the protein failed. This suggests that the glycans are necessary for proper protein processing. Absence of glycans may cause misfolding of the protein during its transport through the secretory pathway causing premature degradation of EXCT4-mgG-2 in this expression system (48).

Due to the high prevalence of HSV-2 infection there is a great need both for a therapeutic and a prophylactic vaccine. Unfortunately, no vaccine has been approved despite great efforts. The major challenge has been that promising results after vaccination in mice and guinea pigs has only in part been translated in clinical trials. For example, Hook et al., showed that the antibody responses to adjuvanted gD-2 in the latest prophylactic vaccine phase III clinical trial HerpeVac (GlaxoSmithKline) demonstrated lack of antibodies identifying three crucial linear epitopes in gD-2 with importance for entry and cell to cell spread, while such antibodies are protective in mice and guinea pigs (49). Although mgG-2 is dispensable for HSV-2 replication in cell cultures, Liljeqvist et. al., demonstrated that mgG-2 negative clinical HSV-2 isolates are rarely detected. Among 2 400 clinical HSV-2 isolates tested using an anti-mgG-2 monoclonal antibody, only two HSV-2 isolates were truly mgG-2 negative, i.e., lack of mgG-2 due to a frameshift mutation and lack of anti-mgG-2 antibodies in patient serum (26). We propose that a prophylactic mgG-2 vaccine against HSV-2 infection may be favourable, not only due to the finding that it is important for preventing neuronal spread, but also because the protein elicits only type-specific HSV-2 immune responses avoiding interference at immunization of cross-reactive immune responses from an earlier HSV-1 infection.

For a therapeutic vaccine in already HSV-2 infected subjects the objective is to reduce clinical lesions and importantly, inhibit or reduce reactivation and thereby preventing both symptomatic and asymptomatic shedding of HSV-2. This task is difficult because the virus has already established latency in sensory ganglia. However, there is an approved vaccine (Shingrix, GlaxoSmithKline) against reactivation of VZV which also establish latency in sensory ganglia. This adjuvanted vaccine use the glycoprotein E produced in CHO cells and prevent herpes zoster in older populations (≥ 50 years of age). The vaccine efficacy was impressively between 96.6% and 97.9% for all age groups (50). This is the first example of a vaccine able to prevent reactivation from the sensory ganglia of a *Alphaherpesvirinae* member. We have recently characterized the antibody responses to EXCT4-mgG-2 antigen in HSV-2 infected patients and showed that the anti-EXCT4-mgG-2 antibodies presented no neutralization capacity but ADCC by human granulocytes, monocytes and natural killer cells, results which are promising for a therapeutic vaccine with the aim to boost the immune responses to mgG-2 in already HSV-2 infected subjects (51).

In conclusion, immunization with recombinant EXCT4-mgG-2 facilitate a glycan dependent protection against neuronal spread after a lethal challenge with HSV-2_WT_, showing that the adaptive immune response can inhibit important functions of mgG-2. Also, we present data showing that mgG-2 is important for the spread of HSV-2 to the neuronal tissue. Altogether, this supports the potential use of recombinantly expressed EXCT4-mgG-2 as a vaccine candidate against HSV-2 infection and indicates that N- and O-linked glycans may be key components in inducing a robust immune response including CD4+ T-cell activation.

## MATERIAL AND METHODS

### Cells and viruses

GMK-AH1 were cultured in Eagle’s minimal essential medium (EMEM, ThermoFisher) supplemented with 2% inactivated FCS and 1% penicillin-streptomycin (PEST). Stock viruses were produced by infecting GMK-AH1 cells without serum in roller bottles at a MOI of 0.5. After complete development of cytopathic effect, the cells and medium were frozen and thawed. The lysate was centrifuged for 10 min at 3000 × g and the supernatant was harvested and kept at − 80 °C. HSV-2 was quantified by a plaque assay on monolayers of GMK-AH1 cells. The titre of the HSV-2_WT_ was 2.0 × 10^7^ PFU/mL, for HSV-2_ΔmgG-2_ 3.4 × 10^6^ PFU/mL, and for HSV-2_rescue_ 4.6 × 10^7^ PFU/mL. The HSV-2_WT_ strain 333 was a kind gift from the University of Pittsburgh 1993. The strain has been passaged <5 times at our laboratory. The GMK-AH1 cells were supplied by Karolinska Institute in Stockholm. The cells were checked for Mycoplasma contamination by real-time PCR and checked for other bacterial and fungal contamination by culturing on agar plates. Contamination checks were performed at the Department of Clinical microbiology at the Sahlgrenska University hospital.

### Production of recombinant mgG-2

The recombinant mgG-2 construct (EXCT4-mgG-2) codes for amino acid A_345_-D_649_, comprising the extracellular region of mgG-2, V_667_-A_670_ from the transmembrane region and A_671_-D_698_, constituting the intracellular region of mgG-2, retrieved from the HSV-2_WT_ strain 333 (Fig. 1A). The protocol is described in detail elsewhere (52). Briefly, the CHO-K1 (028-W4) with the GS expression system was used (Lonza). 50 µM L-methionine sulfoximine (MSX) (Sigma Aldrich) was used as a selection agent for the transfected plasmid. The cells were kept under MSX selection pressure for 30 days before transferring to a 250 mL spinner flask (Bellco) containing 75 mL Dulbecco’s Modified Eagle Medium (L-glutamine-free, Gibco, ThermoFisher Scientific), 25 mL CD FortiCHO (ThermoFisher Scientific) and 50 µM MSX. Protein purification was performed using an ion exchange column (HiPrep QFF 16/10 (Cytiva,)) followed by a HiPrep 16/60 Sephacryl S-400 high resolution gel filtration column (Cytvia).

### Deglycosylation of EXCT4-mgG-2

For complete deglycosylation of EXCT4-mgG-2(−N−O), EXCT4-mgG-2 was incubated with 1× glycobuffer 2, α2-3, 6, 8 neuraminidase (25 U/µg EXCT4-mgG2), O-glycosidase (20 000 U/µg EXCT4-mgG2), α-N-acetyl-galactosaminidase (10 U/µg EXCT4-mgG2) and PNGaseF (250 U/µg EXCT4-mgG2). Removal of only N-linked glycans from (EXCT4-mgG-2(-N) was performed using incubation with 1× glycobuffer 2 and PNGaseF (250 U/µg EXCT4-mgG2). Removal of O-linked glycans from EXCT4-mgG-2(-O) was performed using incubation with 1× glycobuffer 2, α2-3, 6, 8 neuraminidase (25 U/µg EXCT4-mgG2), O-glycosidase (20 000 U/µg EXCT4-mgG2) and α-N-acetyl-galactosaminidase (10 U/µg EXCT4-mgG2). Removal of sialic acids from EXCT4-mgG-2(-SA) was performed using incubation with 1× Glycobuffer 2 and α2-3, 6, and 8 neuraminidases (50 U/µg EXCT4-mgG2). The enzyme control contained 1× Glycobuffer 2, α2-3, 6, 8 neuraminidase (2500 U/mL), O-glycosidase (2 000 000 U/mL), α-N-acetyl-galactosaminidase (1000 U/mL) and PNGaseF (25 000 U/mL). All incubations were performed at 37 °C for 24 hours. All enzymes and glycobuffer 2 were purchased from New England Biolabs. Deglycosylation was confirmed using Western blot and lectin binding assay as described below. To remove free sugar residues, the protein preparations were dialyzed for 24 hours in PBS using the Slide-A-Lyzer™ Mini Dialysis Device 7K MWCO (ThermoFisher Scientific), with medium exchanges at hours 2, 4, 20 and 22.

Deglycosylation of EXCT4-mgG-2 was verified by western blot and lectin blot. In brief, the protein fractions were mixed with 2x SDS-PAGE sample buffer (25% 0.5M Tris-HCl, 20% SDS (10%), 40% glycerol, 10% 2-mercaptoethanol, 5% bromophenol blue) and heated to 95 °C for 10 minutes prior to separation on a NuPage Bis-tris 4-12% gel using MOPS running buffer (Invitrogen) at 100 V for 60 min using an EI9001-XCELL II Mini Cell (Novex) together with a Powerease 500 (Novex). The protein construct was transferred onto an Immobilon PVDF membrane in blotting buffer (NuPAGE™ Transfer Buffer; (ThermoFisher Scientific), 10% methanol). The membrane was blocked in blocking buffer (2% milk, 4% FCS) for 1 hour. As the primary antibody, a monoclonal mouse anti-mgG-2 antibody (17) was used at a 1:2000 dilution, followed by incubation with a horse radish peroxidase-conjugated anti-mouse IgG (Aglient) at a 1:20 000 dilution. Bound antibodies were developed using SuperSignal® West Dura Extended Duration Substrate (ThermoFisher Scientific) and detected in the ChemiDoc™ MP Imaging System (BioRad Laboratories).

Lectin blot was performed by loading, 2 µg each of EXCT4-mgG-2, EXCT4-mgG-2(−N−O), EXCT4-mgG-2(-N), EXCT4-mgG-2(-O) and EXCT4-mgG-2(-SA) to a NuPAGE Bis-Tris 4-12% gel (Invitrogen) and separated using MOPS buffer (Invitrogen) at 200 V. The separated proteins were transferred to a polyvinylidene difluoride membrane (Immobilon-FL 0.45 μm, (Merck Millipore)) using the SemiDry Transblot SD (Bio-Rad Laboratories). Membranes were blocked with block buffer (2% BSA Factor V (Sigma-Aldrich) and 0.1% Tween-20 (VWR chemicals) in PBS (Medicago) followed by incubation with biotinylated lectins; 3 µg/mL Concavalin A (Con A) (Vector Laboratories), 3 µg/mL Jacalin (Vector Laboratories) or 5 µg/mL Maackia Amurensis Lectin II (MAL II, Vector Laboratories) diluted in PBS-BSA-T, at 4 °C for 16-20 hours. Next, membranes were washed three times with 0.1% Tween-20 in PBS (Medicago) followed by incubation with Streptavidin-alkaline phosphatase diluted 1:2000 (Southern Biotech) for 1 h at 20-22 °C. Membranes were washed three times with 0.1% Tween-20 in PBS and developed using BCIP/NBT (Sigma Aldrich) for 1 minute.

### Sample preparation prior to LC-MS/MS

The recombinant EXCT4-mgG-2 samples (24 µg) were constituted in 0.1% sodium deoxycholate (SDOC) (Sigma Aldrich) in 50 mM triethylammonium bicarbonate (Sigma Aldrich) at 0.3 mg/mL. Reduction was done with 5 mM Tris(2-carboxyethyl)phosphine (Sigma Aldrich) at 58 °C for 20 min. The samples were then divided in two parts and digested with either pronase (Roche) or alpha-lytic protease (Sigma Aldrich) for 16 h at 37 °C, at protein:enzyme ratios of 1:20. SDOC was removed by acidification with trifluoroacetic acid followed by centrifugation and then desalted using peptide desalting spin columns (Pierce, ThermoFisher Scientific) according to the manufactureŕs protocol. The eluates were dried in a vacuum centrifuge and dissolved in 3% acetonitrile, 0.1% trifluoroacetic acid (Sigma Adrich) prior to LC-MS/MS analysis.

### LC-MS/MS analysis

Nano-LC-MS/MS was done with the Orbitrap Exploris 480 mass spectrometer (ThermoFisher Scientific) interfaced with an Easy nLC 1200 liquid chromatography system. Analytes were separated using a trap column (15 cm × 0.1 mm, 5 μm particle size Acclaim Pepmap, ThermoFisher Scientific) and an in-house packed C18 analytical column (35 cm × 0.075 mm, 3 μm particle size (Dr. Maisch)) using an acetonitrile gradient in 0.2% formic acid during 90 min. Precursor ion mass spectra were acquired at 120 000 resolution with the scan range m/z 350-2000 and MS/MS analysis with a first mass of m/z 110. Higher-Energy Collisional Dissociation spectra of the most intense precursor ions were performed in data-dependent mode using 3s cycle times, a resolution of 30 000 and a maximum injection time of 100 ms. The normalized automatic gain control target was set to 150 %. Normalized collision energies of 20 % and 30 % were applied on each precursor ion using an isolation window of two mass units. The exclusion time was 12 s. Each sample was run in triplicates using consecutive injections.

### LC-MS/MS Data processing

Data analysis was performed with Proteome Discoverer version 2.4 (ThermoFisher Scientific) using the Byonic (Protein metrics) and Minora (ThermoFisher Scientific) feature detector nodes. The O-glycan modifications were set to ‘6 most common’ with the addition of the HexNAc(2)Hex(2), HexNAc(2)Hex(2)NeuAc(2) and HexNAc(2)Hex(2) NeuAc(4) compositions. The number of allowed O-glycans per peptide was 0-2 of the same or with different compositions. The N-glycan modifications were set to ‘57 human plasmà. Additional allowed modifications were methionine oxidation and the Asn to Asp modification. Free cleavage sites were searched using the mgG-2 protein construct sequence, at a 5 ppm mass limit for the precursor ions and a 20 ppm mass limit for the MS2 ions. All glycopeptide identities were manually verified to contain the presence of the correct peptide and/or peptide+HexNAc ion and the proper set of oxonium ions. For instance, presence of the oxonium ion at m/z 274.09 for compositions including Neu5Ac, presence of m/z 366.14 for compositions including HexHexNAc and m/z 163.06 for compositions including oligomannose N-glycosylation. The maximum intensity measurement by Minora was used for the relative quantification measurements. In addition, manual inspection of the raw spectra using Xcalibur qual browser (ThermoFisher Scientific) was performed to confirm the presence of the N-linked glycoforms. The CVs for all glycoforms are based on the percental distribution within each LC-MS/MS injection, glycoforms with a relative abundance < 1% are excluded from the calculations.

Of note, the m/z value of the oxonium ions obtained during LC-MS/MS analysis is 204 for GlcNAc and GalNAc alike. Similarly, the addition of a Gal or Glc on the N-acetylated hexose (e.i GlcNAc or GalNAc) results in a peak at m/z 366 in the MS2 spectra. Hence, in our assay, it was not possible to distinguish between the types of hexoses. However, due to extensive previous glycoproteomic analysis of the HSV-2 and CHO-cell lines, done by us and others, and the confirmation of O-linked core 1 glycan structures, the m/z 204 and 366 peaks most likely correspond to GalNAc and GalGalNAc, respectively.

### Mouse model for immunization and HSV-2 genital challenge

The animals were kept at Experimental Biomedicine, Gothenburg University, and the protocol was approved by the local ethics committee, and all procedures were performed according to approved guidelines and regulations. Female 6- to 8-week-old C57BL/6 mice were intramuscularly immunized three times at 10-day intervals using 2.5 µg EXCT4-mgG-2, EXCT4-mgG-2(−N−O), EXCT4-mgG-2(-N), EXCT4-mgG-2(-O), EXCT4-mgG-2(-SA) or controls (PBS alone). In all experiments adjuvant consisting of 250 µg alum (Alhydrogel) + 20 µg CpG (ODN1826, TCC ATG ACG TTC CTG ACG, TT), (Operon Biotechnologies GmbH) were used and diluted in 30 µL Tris-buffered saline (TBS). Six days prior to challenge, the mice were injected subcutaneously with 3 mg Depo-Provera (medroxyprogesterone acetate) (Pfizer). The mice were challenged intravaginally with 1 × 10^5^ PFU of HSV-2_WT_ (corresponding to 25 × LD_50_). Mice were scored daily for vaginal inflammation and disease and were graded as follows: 0 - no symptoms (healthy); 1 - genital erythema; 2 - moderate genital inflammation (blisters); 3 - severe symptoms or neurologic symptoms (purulent genital lesions, loss of hair); 4 - paralysis or generally poor condition. Mice presenting a score of 3 or higher were euthanized.

### Quantification of total IgG in mouse serum

Blood samples were drawn from the tail vein 14 days after the third immunization. Sera were separated from whole blood using centrifugation twice for 15 minutes at 400 g. ELISA plates (96-well) were coated with 100 µL (1 µg/mL diluted in carbonate buffer (pH 9.6)) EXCT4-mgG-2, EXCT4-mgG-2(−N−O), EXCT4-mgG-2(-N), EXCT4-mgG-2(-O) or EXCT4-mgG-2(-SA), for serum samples collected from mice immunized with EXCT4-mgG-2 (n=29), EXCT4-mgG-2(−N−O) (n=16), EXCT4-mgG-2(-N) (n=10), EXCT4-mgG-2(-O) (n=10) or EXCT4-mgG-2(-SA) (n=10), respectively. Plates were blocked with 2% milk, and 100 µL of serum samples diluted 1:90, 1:270, 1:810, 1:2430, 1:7290, 1:21870, 1:65610, and 1:196830 in 1% milk + 0.05% Tween in PBS were added to the wells. The plates were incubated for 1.5 hours at 37 °C before washing with TBS that contained 0.05% Tween, followed by the addition of peroxidase-conjugated anti-mouse IgG (Sigma Aldrich) and incubation for 1.5 hours at 37 °C. Peroxidase activity was detected by the addition of o-phenylenediamine dihydrochloride (Sigma Aldrich), in citrate buffer. The reaction was stopped with sulfuric acid after 30 seconds, and the OD was measured at 490 nm in a microplate reader. The antibody titre was defined as the highest serum dilution giving an OD ≥ 0,250, defined as OD 0,2 above the value of the negative control.

### Serum IgG reactivity against recombinant mgG-2

96-well Maxisorp plates were coated with EXCT4-mgG-2 or EXCT4-mgG-2(−N−O) (1 µg/mL diluted in carbonate buffer (pH 9.6)) and incubated at 4 °C, overnight. Blocking of the plates was performed with 2 % milk in PBS for 30 minutes at room temperature. Serum samples from mice immunized with EXCT4-mgG-2 or EXCT4-mgG-2(−N−O) was diluted 1:100 in dilution buffer (1 % milk and 0.05 % tween in PBS) and incubated for 1.5 h at room temperature followed by washing of the plates and addition of Alkaline phosphatase-conjugated anti-mouse IgG (Jackson ImmunoResearch Laboratories) diluted 1:1000 in dilution buffer. Following 1.5 hours incubation at room temperature the plates were again washed and 1 mg/mL p-nitrophenylphosphate (Medicago) dissolved in diethanolamine buffer was added. The plates were incubated 15 minutes in the dark before OD measurement at 405 nm in a microplate reader.

### Viral plaque assay

To measure viral replication in the genital tract, vaginal washes were collected from immunized and infected mice 2-days post-infection. From mice infected with the HSV-2_WT_, HSV-2_ΔmgG-2_ or HSV-2_rescue_ (see below) vaginal fluids were collected at 12 h and at 1-, 2-, 3-, 4-, 5- and 6-days after infection. Washes were obtained by gently pipetting 40 μl of Hanks balanced salt solution (HBSS) in and out of the vagina until a clump of mucus was retrieved, followed by a second wash with an additional 40 μl. Both washes were pooled and collected in a total volume of 1 ml HBSS. Samples were stored at − 80 °C until analysis. Also, the vaginas from mice infected with HSV-2_WT_, HSV-2_ΔmgG-2_ or HSV-2_rescue_ was excised and homogenized (see section Tissue sample collection for HSV-2 DNA quantification below) and assessed for infectious viral particles using a plaque assay. To perform the plaque assay, samples were diluted 1:10 in

PBS and added to 80 % confluent monolayers of GMK-AH1 cells grown in 6- or 24-well plates, after which the plate was incubated at room temperature for 1 hour. Methylcellulose was mixed with EMEM (ThermoFisher Scientific) supplemented with 1% PEST and 2% FCS, and a total of 2 mL (6-well plates) or 400 µL (24-well plates) were added on top of the cultured cell layer. The plates were incubated for 72 h at 37 °C and 4% CO_2_ before removal of the methylcellulose overlay and staining with one drop of crystal violet per well. Thereafter the cells were washed with tap water four times and allowed to dry. The plaques were counted using a light microscope at four times magnification.

### HSV-2 DNA quantification in tissue samples

The DRG and the spinal cord were collected from mice immunized with EXCT4-mgG-2, EXCT4-mgG-2(−N−O), EXCT4-mgG-2(-N), EXCT4-mgG-2(-O), or EXCT4-mgG-2(-SA), either when the mice presented a disease score ≥ 3, or upon termination of the experiment at day 14. Samples were stored at -80 °C until further analysis. Ganglia and spinal cord were separately transferred to MagNa Lyser Green Bead tubes (Roche) and mixed with lysis buffer from the MagNa Pure LC DNA Isolation Kit II Tissue (Roche). The tissues were homogenized twice for 54 s at 6500 rpm in a MagNa Lyser (Roche). Viral DNA was extracted on a MagNa Pure LC Robot (Roche) together with the MagNA Pure LC DNA Isolation Kit I for DNA from serum or vaginal washes, or with the MagNA Pure LC DNA Isolation Kit II Tissue for DNA from tissue samples, according to the instructions from the manufacturer. Isolated DNA was stored at -80 °C until further analysis.

Quantification of viral DNA was performed with the 7300 Real-Time PCR System (Applied Biosystems) in a 50 µL reaction that contained 10 µg extracted DNA Universal Mastermix (Thermo Fisher), and a TaqMan probe and primers with the following sequences; forward primer, 5’-TGC AGT TTA CGT ATA ACC ACA TAC AGC; reverse primer, 5’-AGC TTG CGG GCC TCG TT; and the probe, 5’-CGC CCC AGC ATG TCG TTC ACG-VIC-TAMRA (53). For standardization of HSV-2 DNA to cell content a beta-globin gene system was constructed from a conserved region of the gene. The primers were as follows: forward; 5’-CTG AAA CAC TAT GGT GGA GCT CA, and reverse, 5’-AAC ACC AAG TTC TTC TGC CTT CAC. The probe, sequence 5’-TGC AGA GGA GAA GGC AGC CAT CAC T-FAM-BHQ1 (black hole quencher 1). The PCR reaction was run for 2 minutes at 50 °C, 10 minutes at 95 °C, followed by 45 cycles of 15 seconds at 95 °C and 1 minute at 58 °C.

In addition, Plasmids (pUC57) containing the target sequences were constructed (GenScript) and amplified in E. coli XL-1 Blue, purified by HiSpeed Plasmid Maxi Kit 7 (Qiagen) and quantified by spectrophotometer analysis. A standard curve based on six five-fold dilutions of the plasmid using an initial concentration of 1 × 10^6^ HSV-2 DNA and beta-globin copy numbers per reaction was included in each run. For tissue samples from lymph nodes and spinal cord we observed an inhibition of the PCR reaction probably due to high content of cellular DNA. These samples were therefore diluted 1:4 in PBS before extraction of HSV-2 DNA.

### Mouse model to assess T-cell specific immune responses

The vaccine candidates EXCT4-mgG-2 and EXCT4-mgG2(−N−O) were tested in 8-week-old female C57BL/6 mice (Charles River). Animals were kept at the Astrid Fagraeus laboratory, Karolinska Institutet, in accordance with the recommendations of the Swedish Board of Agriculture. The protocol was approved by the local ethics committee, and all animal procedures were performed according to the approved guidelines and regulations. Protein antigen preparations were formulated in CpG1826 and alum (both from Invivogen) and inoculated intramuscularly in the gastrocnemius muscle of the left hind leg, three times at 10-day intervals at the doses of 2.5 μg antigen per mouse.

### Isolation of mouse splenocytes

Mouse spleens were harvested 14-days post the last immunization and mashed through 70-μm-pore-size nylon cell strainers to obtain single-cell suspensions. Cells were washed in complete RPMI medium (RPMI 1640 supplemented with 5% FCS, 2 mM l-glutamine, 100 U/ml penicillin, and 100 μg/ml streptomycin (all from Gibco/Invitrogen), followed by treatment with red blood cell lysis buffer (Sigma Aldrich) for 2 min. After another wash, the cells were resuspended in complete RPMI medium. Viable cells were quantified by using Guava Viacount and Guava cell counter (Merck-Millipore).

### FluoroSpot analysis

Mouse IFN-γ/IL-2/TNF FluoroSpot assay were performed using a commercial kit (Mabtech) according to the manufacturer’s instructions. Briefly, pre-coated plates for IFN-γ, IL-2, and TNF were washed five times with PBS. Splenocytes were plated at the concentration of 200 000 cells per well and stimulated with EXCT4-mgG-2 antigen 4µg/ml, EXCT4 peptide pool 4µg/ml (GenScript), ConA 4µg/ml (Merck-Millipore) or complete RPMI medium, then incubated for approximately 42 h at 37 °C in 5% CO₂. After incubation, cells were removed, and plates were washed before the addition of fluorophore-conjugated detection antibodies. Following further washing and drying, spots were visualized and quantified using a FluoroSpot reader (Mabtech IRIS).

### Intracellular cytokine assay

Freshly isolated splenocytes were set up at 2 × 10^6^ splenocytes per well in complete RPMI medium in 96-well round-bottom plate. Samples were incubated overnight at 37 °C in 5% CO₂ with 2 µg/ml EXCT4-mgG-2 peptide pool or complete medium alone. Cells were incubated with Protein Transport Inhibitor Cocktail (eBiosciences) for 4 hours. Cells were washed with FACS buffer and incubated with anti-CD3e-PerCP-Cy5.5 (145-2C11), anti-CD4-FITC (GK1.5), and anti-CD8-APC (53-6.7) in FACS buffer (all monoclonal antibodies were purchased from eBiosciences). After the excess antibody was washed away, the cells were fixed with Intracellular Fixaton & Permeabilization Buffer (eBiosciences) according to the manufacturer’s instructions. Samples were then stained with anti-IFN-γ-PE (XMG1.2) (eBiosciences). After being washed the cells were resuspended in FACS buffer and analysed with the Guava cell counter (Merck-Millipore).

### Frame shift mgG-2 negative HSV-2 mutant and rescue

An mgG-2 negative mutant was selected from the wild type HSV-2 strain 333 (referred in this study as HSV-2_WT_ (Fig. 1B)) in GMK-AH1 cells in the presence of a sulphated oligosaccharide. This mutant, designated as AC9 (accession number EU018128), presented a deletion of a single nucleotide C (frame shift mutation) within a run of seven cytosine nucleotides (nts) at 1649-1655 resulting in a premature termination codon (TAA) at nts 1924-1926. A PCR amplified fragment encompassing nts 222-2140 of the gG-2 gene from strain AC9 was introduced back to HSV-2_WT_ by a marker transfer assay. This virus is designated in this study as HSV-2_ΔmgG-2_ (25), (Fig. 1C). To rescue the HSV-2_ΔmgG-2_ mutant the gG-2 gene fragment (nts 222-2140) from the HSV-2_WT_ genome was amplified by PCR and co-transfected together with HSV-2_ΔmgG-2_ genomic DNA into GMK-AH1 cells. The selection of positive variants was based on a plaque assay using an anti-mgG-2 MAb (O1.C5.B2) (18) followed by the amplification of the gG-2 gene and DNA sequencing to confirm the wild type sequence. The marker rescued HSV-2 is designated in this study as HSV-2_rescue_. Primers for sequencing of the gG-2 gene were used as described (26).

As shown in the schematic illustration of the gG-2 proteins for strain HSV-2_ΔmgG-2_ (Fig. 1C) the mutation disrupts the protein sequence of mgG-2 from amino acid 552 including the immunodominant region (amino acids 551-573) (14), the transmembrane region as well as the intracellular region. The HSV-2_ΔmgG-2_ showed to lack expression of mgG-2 (25, 29). In addition, as the viral gG and Us3 genes are expressed from the same bicistronic gene the normal expression of the Us3 protein was confirmed earlier (29). To investigate that sgG-2 is not affected by the frame shift mutation in mgG-2 we infected GMK-AH1 cells at a MOI of 1 using HSV-2_ΔmgG-2_, HSV-2_rescue_ and HSV-2_WT_. After 24-36 h, at complete cytopathic effect, cell culture medium was centrifuged at 4000 × g. As the amount of PFU at complete cytopathic effect is different for the three strains we normalized the amount of supernatant to equal PFU with the ratio 1.0 for HSV-2_ΔmgG-2_ (titre 3.4 × 10^6^)_,_ ratio 0.17 for the HSV-2_WT_ (titre 2.0 × 10^7^), and ratio 0.074 for the HSV-2_rescue_ strain (titre 4.6 × 10^7^). The supernatant was mixed with sample buffer containing SDS and mercaptoethanol, followed by boiling 10 min and separated on a 4-12% NuPAGE Bis-Tris gradient gel (Invitrogen) with 3-(N-morpholino) propane sulfonic acid and SDS as running buffer. The proteins were transferred to an immobilon-PVDF membrane. A pre-stained protein ladder (ThermoFisher Scientific) was used as molecular weight marker. The anti-sgG-2 MAb (4.A5.A9) (17) was used for detection, at a 1:2000 dilution, and HRP-conjugated polyclonal anti-mouse IgG was used as conjugate (DakoCytomation), at a 1:20 000 dilution. After washing, SuperSignal® West Dura Extended Duration Substrate (ThermoFisher Scientific) was added followed by detection using the ChemiDoc™ MP Imaging System (BioRad Laboratories).

### Mouse model of genital HSV-2 infection

Female 6- to 8-week-old C57BL/6 mice (Scanbur BK) were used. The animals were kept at Experimental Biomedicine, Gothenburg University, and the protocol was approved by the local ethics committee, and all procedures were performed according to approved guidelines and regulations. The mice were anesthetized with 3 % isoflurane (Baxter). Before infection 3 mg of Depo-Provera, was given subcutaneously and 6 days later 4 × 10^4^ PFU (corresponding to 10 × lethal dose 50% (LD_50_)) of HSV-2_WT_, HSV-2_ΔmgG-2_ or HSV-2_rescue_ were given intravaginally with a blunted fine plastic pipette. We also used the more sensitive mouse strain DBA/2 (Harlan Lab) and infected 6- to 8-weeks old female mice intravaginally with 1.25 × 10^4^ PFU (corresponding to 125 × LD_50_) of HSV-2_WT_, HSV-2_ΔmgG-2_ or HSV-2_rescue_. The natural course of the vaginal HSV-2 infection is described in detail by Parr and Parr (28). Briefly, HSV-2 infects initially the vaginal epithelium followed by spread to vulva, anus, and perineum, and to the autonomic and to the sensory neurons in DRG and the spinal cord. Mice were scored as described above.

### Tissue sample collection for HSV-2 DNA quantification

For the C57BL/6 mice, the vagina was excised at 12 h and at 1-, 2-, 3-, 4-, 5- and 6-days after infection and kept in 1 mL PBS at -80 °C. DRG and spinal cord were collected at 3- or 6-days after infection with HSV-2_WT_ or HSV-2_Δmg-G_ and at 14- or 21-days after infection of C57BL/6 mice infected with HSV-2_Δmg-G_. From C57BL/6 mice, serum was drawn daily from day 1 to day 6 and inguinal lymph nodes were collected at day 1 or day 6. From DBA/2 mice serum and inguinal lymph nodes were collected at day 6 post infection. All tissue samples were kept at -80 °C, thawed and homogenized by using a Dounce homogenizer prior to DNA extraction using the MagnaPure LC Robot (Roche) and DNA quantification using qPCR as described above.

### Tissue preparation, histology, and immunohistochemistry

DBA/2 mice were infected intravaginally with 1.25 x10^4^ PFU of HSV-2_ΔmgG-2_ or HSV-2_rescue_. At day 6 the animals were deeply anesthetized with isoflurane and fixed by transcardial perfusion via the left ventricle with PBS, followed by 4 % formaldehyde. The spinal cord including DRG was removed, and immersion fixed for about 4 days. After fixation the tissue was decalcified, using 10 % EDTA in 0.2 M Tris buffer, pH 7.4 for 3-4 weeks; the buffer was changed twice weekly. After decalcification, 3-4 mm thick horizontal slices were cut from the lumbosacral, the lumbar and thoracic level of the spinal cord. The specimens were dehydrated, infiltrated, and embedded in paraffin using a Leica TP1020 Automatic Tissue Processer (Leica Biosystems). Sections were cut at 4 μm and, after high temperature antigen retrieval, using a 0.01 M citrate buffer, pH 6.0, sections were incubated with rabbit polyclonal antibodies against HSV-2 antigen diluted 1/4000 (DakoCytomation). Anti-rabbit Impress Reagent HRP (Vector laboratories) was used as secondary reagents, and the immunoreactions were visualized using liquid DAB+ substrate (DakoCytomation). Nuclei were counter-stained with haematoxylin. Finally, the sections were dehydrated and mounted using DPX (Merck). A Zeiss Axio Image M1 microscope and AxioVision software (Zeiss) was used for documentation.

### Ethics

The animal studies were approved by the Swedish board of agriculture (Dnr 5.8.28-18516/2018), (Dnr 5.8.18-08670/2022) and (Dnr 6054-2021).

### Statistics

The SPSS 16 (IBM) and the GraphPad Prism 10.4.1 (Dotmatics) software programs were used for statistical calculations. Mann-Whitney or the Kruskal-Wallis (non-parametric) tests were used for comparison of HSV-2 viral load (PFU and HSV-2 DNA genome copies), and for comparison of IgG levels. The area under curve method based on the trapezoidal rule included in SigmaPlot was used for calculation of the ratio of PFU in vagina and vaginal washes. For survival, the pairwise Log-rank (Mantel-Cox) test was used to test significantly altered survival as compared to mice immunized with EXCT4-mgG-2 or infected with HSV-2_wt_. For the fluorospot and intracellular cytokine assays, the data passed as normally distributed using the Shapiro-Wilk test, and significance was tested using the Tukey’s multiple comparison test. P-values < 0.05 were considered statistically significant:

## ACKNOWLEDGEMENTS

Proteomic analysis was performed at the Proteomics Core Facility, Sahlgrenska academy, Gothenburg University, with financial support from SciLifeLab and BioMS. We acknowledge the Mammalian Protein Expression Core Facility at the University of Gothenburg for technical support in lectin blotting and recombinant protein production.

## FUNDING

The work was supported by grants from Sweden’s innovation agency Vinnova (2020–03108) and the ALF Foundation of Sahlgrenska University Hospital (ALFGbg-1006865 and ALFGbg-716041).

## CONFLICT OF INTEREST

J.L., L.G., S.G., and T.B. develop a vaccine against HSV-2 infection in the company Simplexia AB. No financial contributions to this work have been received from Simplexia AB. All other authors report no conflicts of interest.

## AUTHOR CONTRIBUTIONS

Conceptualization, E.K., J.L. and R.N.; methodology, E.K., C.G., L.G., J.N., M.E., B.A., S.G., E.J., S.L, and E.T.; data curation, validation, and analysis, E.K., L.G., E.M., J.N. and S.G., E.T. ; resources, E.M., L.G., J.L., and R.N.; writing — original draft preparation, E.K., J.L. and R.N.; writing — review and editing, E.K., L.G., E.M., J.N., S.G., E.T., T.B., J.L. and R.N. visualization, E.K., J.N., S.G., E.J., and S.L.; supervision, E.M., J.L., and R.N.; funding acquisition, J.L., and R.N. All authors have read and agreed to the published version of the manuscript.

