## Supplemental material for "Glycoprotein G enables HSV-2 neuroinvasion and provides protection as a glycosylated vaccine antigen"

**SUPPLEMENTARY MATERIAL**

**Table S1.** Observed O-glycopeptides in the EXCT4-mgG-2 preparation. CV is calculated based on the percentual distribution within each injection (n = 3). For peptides with multiple possible glycan sites, one glycan structure was identified but the exact position cannot be determined.

| Glycan site | Peptide Sequence | Glycan Composition | Theo. MH+ [Da] | Precursor ion Abundances (AU, Average for three injections) | Abundance (%, average for three injections) | Abundance CV (%) | Assigned glycan type |
|---|---|---|---|---|---|---|---|
| T356 | [L].TEDASSDSPTSAPEKTPLPV.[S] | NG | 2028,961 | 3,66E+06 | 5,72 | 6,71 | |
| T356 | [L].TEDASSDSPTSAPEKTPLPV.[S] | HexNAc(1)Hex(1) | 2394,093 | 9,77E+06 | 15,27 | 10,38 | O-linked |
| T356 | [L].TEDASSDSPTSAPEKTPLPV.[S] | HexNAc(1)Hex(1) NeuAc(1) | 2685,188 | 3,23E+07 | 50,63 | 5,03 | O-linked |
| T356 | [L].TEDASSDSPTSAPEKTPLPV.[S] | HexNAc(1)Hex(1) NeuAc(2) | 2976,284 | 1,81E+07 | 28,38 | 3,55 | O-linked |
| S411/T415 | [A].VASPPATA.[S] | NG | 713,3828 | 1,71E+06 | 2,41 | 23,71 | |
| S411/T415 | [A].VASPPATA.[S] | HexNAc(1) | 916,4622 | 3,55E+06 | 4,97 | 9,66 | O-linked |
| S411/T415 | [A].VASPPATA.[S] | HexNAc(1)Hex(1) | 1078,515 | 4,22E+06 | 5,90 | 7,55 | O-linked |
| S411/T415 | [A].VASPPATA.[S] | HexNAc(1)Hex(1) NeuAc(1) | 1369,61 | 4,29E+07 | 59,47 | 1,74 | O-linked |
| S411/T415 | [A].VASPPATA.[S] | HexNAc(2)Hex(2) | 1443,647 | 2,92E+06 | 4,07 | 6,44 | 2 x O-linked |
| S411/T415 | [A].VASPPATA.[S] | HexNAc(1)Hex(1) NeuAc(1); HexNAc(1)Hex(1) | 1734,743 | 9,05E+06 | 12,56 | 4,93 | 2 x O-linked |
| S411/T415 | [A].VASPPATA.[S] | HexNAc(2)Hex(2) NeuAc(2) | 2025,838 | 7,68E+06 | 10,63 | 3,59 | 2 x O-linked |
| S417/S420/S421 | [A].SVESSPLPA.[S] | NG | 886,4516 | 1,04E+07 | 22,50 | 6,60 | |
| S417/S420/S421 | [A].SVESSPLPA.[S] | HexNAc(1) | 1089,531 | 2,80E+07 | 7,88 | 17,54 | O-linked |
| S417/S420/S421 | [A].SVESSPLPA.[S] | HexNAc(1)Hex(1) | 1251,584 | 2,65E+07 | 1,12 | 16,57 | O-linked |
| S417/S420/S421 | [A].SVESSPLPA.[S] | HexNAc(1)Hex(1) NeuAc(1) | 1542,679 | 9,09E+07 | 2,83 | 56,86 | O-linked |
| S417/S420/S421 | [A].SVESSPLPA.[S] | HexNAc(1)Hex(1) NeuAc(2) | 1833,775 | 3,91E+06 | 13,26 | 2,44 | O-linked |
| T445/T448/T449/T453/T454 | [A].AKTPPTTPAPTTPPPTST.[H] | NG | 1762,922 | 8,51E+06 | 86,66 | 0,73 | |

| | | | | | | | |
|---|---|---|---|---|---|---|---|
| T445/T448/T449/T453/T454 | [A].AKTPPTTPAPTTPPPTST.[H] | HexNac(1)Hex(1) | 1966,002 | 1,31E+06 | 13,34 | 4,77 | O-linked |
| T529/T532/T533 | [R].TPPTDPKTHPHGPA.[D] | NG | 1452,723 | 5,44E+08 | 31,03 | 5,39 | |
| T529/T532/T533 | [R].TPPTDPKTHPHGPA.[D] | HexNAc(1) | 1655,802 | 2,21E+08 | 12,67 | 6,44 | O-linked |
| T529/T532/T533 | [R].TPPTDPKTHPHGPA.[D] | HexNAc(1)Hex(1) | 1817,855 | 2,49E+08 | 14,16 | 5,42 | O-linked |
| T529/T532/T533 | [R].TPPTDPKTHPHGPA.[D] | HexNAc(2) | 1858,882 | 7,50E+07 | 4,28 | 4,87 | 2 x O-linked |
| T529/T532/T533 | [R].TPPTDPKTHPHGPA.[D] | HexNAc(2)Hex(1) | 2020,935 | 2,00E+08 | 11,36 | 15,60 | 2 x O-linked |
| T529/T532/T533 | [R].TPPTDPKTHPHGPA.[D] | HexNAc(1)Hex(1) NeuAc(1) | 2108,951 | 3,63E+07 | 2,08 | 10,74 | O-linked |
| T529/T532/T533 | [R].TPPTDPKTHPHGPA.[D] | HexNAc(2)Hex(2) | 2182,987 | 2,72E+08 | 15,52 | 10,42 | 2 x O-linked |
| T529/T532/T533 | [R].TPPTDPKTHPHGPA.[D] | HexNAc(2)Hex(2) NeuAc(1) | 2474,083 | 1,29E+08 | 7,33 | 11,51 | 2 x O-linked |
| T529/T532/T533 | [R].TPPTDPKTHPHGPA.[D] | HexNAc(2)Hex(2) NeuAc(2) | 2765,178 | 2,05E+07 | 1,16 | 14,95 | 2 x O-linked |
| S548 | [A].DAPPGSPAPPPPEHR.[G] | NG | 1521,744 | 3,26E+07 | 2,02 | 8,03 | |
| S548 | [A].DAPPGSPAPPPPEHR.[G] | HexNAc(1) | 1724,824 | 8,73E+07 | 5,37 | 6,90 | O-linked |
| S548 | [A].DAPPGSPAPPPPEHR.[G] | HexNAc(1)Hex(1) | 1886,877 | 4,19E+08 | 25,80 | 1,05 | O-linked |
| S548 | [A].DAPPGSPAPPPPEHR.[G] | HexNAc(1)Hex(1) NeuAc(1) | 2177,972 | 9,58E+08 | 59,01 | 0,51 | O-linked |
| S548 | [A].DAPPGSPAPPPPEHR.[G] | HexNAc(1)Hex(1) NeuAc(2) | 2469,068 | 1,27E+08 | 7,80 | 2,68 | O-linked |
| T615 | [L].GPLAPNTPRPPA.[Q] | NG | 1187,653 | 2,81E+06 | 1,95 | 8,16 | |
| T615 | [L].GPLAPNTPRPPA.[Q] | HexNAc(1) | 1390,733 | 6,30E+06 | 4,40 | 8,22 | O-linked |
| T615 | [L].GPLAPNTPRPPA.[Q] | HexNAc(1)Hex(1) | 1552,785 | 4,38E+07 | 30,55 | 4,42 | O-linked |
| T615 | [L].GPLAPNTPRPPA.[Q] | HexNAc(1)Hex(1) NeuAc(1) | 1843,881 | 6,46E+07 | 45,11 | 2,88 | O-linked |
| T615 | [L].GPLAPNTPRPPA.[Q] | HexNAc(1)Hex(1) NeuAc(2) | 2134,976 | 2,58E+07 | 17,98 | 0,46 | O-linked |
| S629/T632 | [A].KDMPSGPTPQHIPL.[F] | | 1517,778 | 7,84E+06 | 1,32 | 2,09 | O-linked |
| S629/T632 | [A].KDMPSGPTPQHIPL.[F] | HexNAc(1)Hex(1) | 1882,91 | 2,81E+07 | 4,72 | 2,13 | O-linked |
| S629/T632 | [A].KDMPSGPTPQHIPL.[F] | HexNAc(1)Hex(1) NeuAc(1); HexNAc(1)Hex(1) NeuAc(2) | 3121,329 | 1,35E+07 | 2,28 | 5,37 | O-linked |
| S629/T632 | [A].KDMPSGPTPQHIPL.[F] | HexNAc(1)Hex(1) NeuAc(1) | 2174,006 | 2,15E+08 | 36,21 | 1,48 | O-linked |
| S629/T632 | [A].KDMPSGPTPQHIPL.[F] | HexNAc(1)Hex(1) NeuAc(2) | 2465,101 | 3,04E+08 | 51,13 | 0,77 | O-linked |
| S629/T632 | [A].KDMPSGPTPQHIPL.[F] | HexNAc(2)Hex(2) NeuAc(2) | 2830,233 | 2,17E+07 | 3,65 | 2,78 | 2 x O-linked |
| S643/S645 | [W].FLTASPALD.[V] | NG | 934,488 | 3,19E+06 | 14,15 | 10,98 | |
| S643/S645 | [W].FLTASPALD.[V] | HexNAc(1)Hex(1) NeuAc(1) | 1590,716 | 8,39E+06 | 27,95 | 6,71 | O-linked |
| S643/S645 | [W].FLTASPALD.[V] | HexNAc(1)Hex(1) NeuAc(2) | 1881,811 | 1,09E+07 | 48,50 | 1,03 | O-linked |

NG = Non glycosylated

**Table S2.** Observed N-glycopeptides in the EXCT4-mgG-2 preparation. CV is calculated based on the percentual distribution within each injection (n = 3).

| Glycan site | Peptide Sequence | Glycan Composition | Theo. MH+ [Da] | Precursor ion Abundances (AU, Average for three injections) | Abundance (%, average for three injections) | Abundance CV (%) | Assigned glycan type | Abundance (%) of complex type structures |
|---|---|---|---|---|---|---|---|---|
| N436 | [A].AAATPGAGHT NTS.[S] | NG | 1155,54 | 3,78E+05 | 4,06 | 138,81 [a] | | |
| N436 | [A].AAATPGAGHT NTS.[S] | HexNAc(2)Hex(3) Fuc(1) | 2193,91 | 3,25E+06 | 28,12 | 12,60 | N-linked, paucimannose | |
| N436 | [A].AAATPGAGHT NTS.[S] | HexNAc(2)Hex(3) Fuc(1); HexNAc(1) | 2396,99 | 2,92E+05 | 2,56 | 5,80 | N-linked, Complex [b] | |
| N436 | [A].AAATPGAGHT NTS.[S] | HexNAc(2)Hex(3) Fuc(1); HexNAc(1)Hex(1) | 2559,05 | 2,64E+05 | 2,31 | 12,65 | N-linked, Complex [b] | |
| N436 | [A].AAATPGAGHT NTS.[S] | HexNAc(2)Hex(4) | 2209,91 | 1,11E+06 | 9,77 | 2,78 | N-linked, Oligomannose | |
| N436 | [A].AAATPGAGHT NTS.[S] | HexNAc(2)Hex(5) | 2371,96 | 2,34E+06 | 20,81 | 8,50 | N-linked, Oligomannose | |
| N436 | [A].AAATPGAGHT NTS.[S] | HexNAc(4)Hex(3) Fuc(1) | 2600,07 | 5,40E+05 | 4,73 | 10,50 | N-linked, Complex | 14,49 |
| N436 | [A].AAATPGAGHT NTS.[S] | HexNAc(4)Hex(4) Fuc(1) | 2762,13 | 1,05E+06 | 9,15 | 10,38 | N-linked, Complex | 28,05 |
| N436 | [A].AAATPGAGHT NTS.[S] | HexNAc(4)Hex(5) Fuc(1) | 2924,18 | 1,57E+06 | 13,64 | 9,46 | N-linked, Complex | 42,05 |
| N436 | [A].AAATPGAGHT NTS.[S] | HexNAc(4)Hex(5) Fuc(1)NeuAc(1) | 3215,27 | 5,75E+05 | 4,87 | 30,60 | N-linked, Complex | 15,42 |
| N511 | [S].AANVSVA.[A] | HexNAc(2)Hex(5) | 1847,76 | 6,67E+06 | 100 | 0 | N-linked, Oligomannose | |

[a] The high CV among this peptide originates from a different peak-integration between injections. However, manual inspection of the peaks could confirm similar elution profiles and intensities in all three injections.
[b] Manual inspection of the ion spectra suggests a monoanntennary N-linked complex-type glycan. NG = Non glycosylated

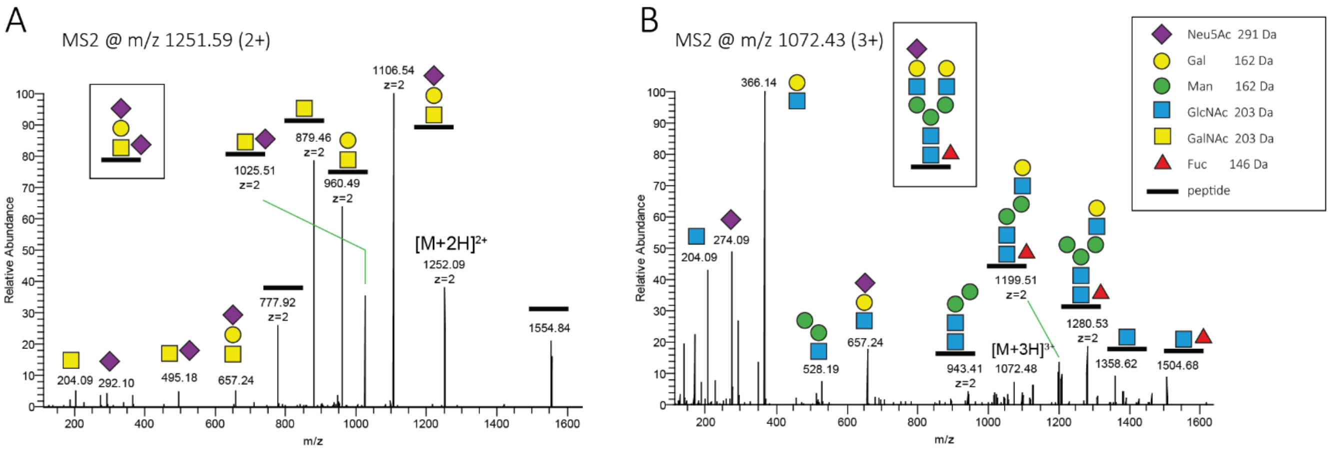


**Figure S1**. Representative MS2 spectra of EXCT4-mgG-2 glycopeptides. **(A)** Fragmentation analysis of [L].GPLAPNTPRPPA.[Q] containing a di-sialylated core 1 O-glycan at T615. **(B)** Fragmentation analysis of [A].AAATPGAGHTNTS.[S] containing a mono-sialylated and core-fucosylated complex biantennary N-glycan at N436. The precursor ion structures are shown in the boxes. Ion charges are indicated when z>1.

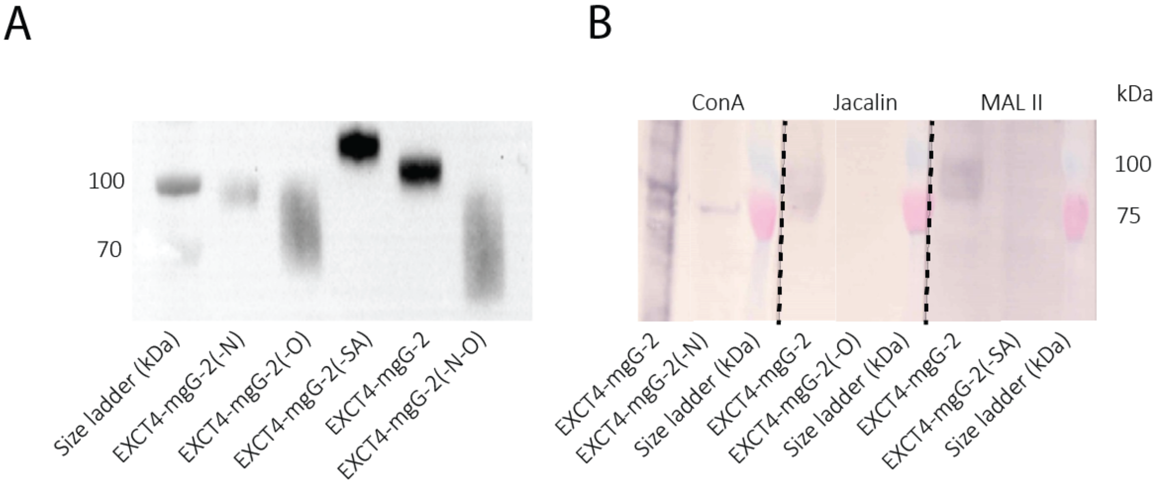


**Figure S2.** Verification of the enzymatic removal of glycan structures. **(A)** Western blot showing size shift of the N- and O-glycosylated EXCT4-mgG-2 and of the deglycosylated EXCT4-mgG-2(-N), EXCT4-mgG-2(-O), EXCT4-mgG-2(-SA), and EXCT4-mgG-2(−N−O). **(B)** Lectin blot confirming the removal of specific glycan structures. ConA; Binds to core oligomannose of N-linked glycans, Jacalin; binds to Tn and sialylated Tn-antigens of O-linked glycans, MAL II; binds to sialic acids.

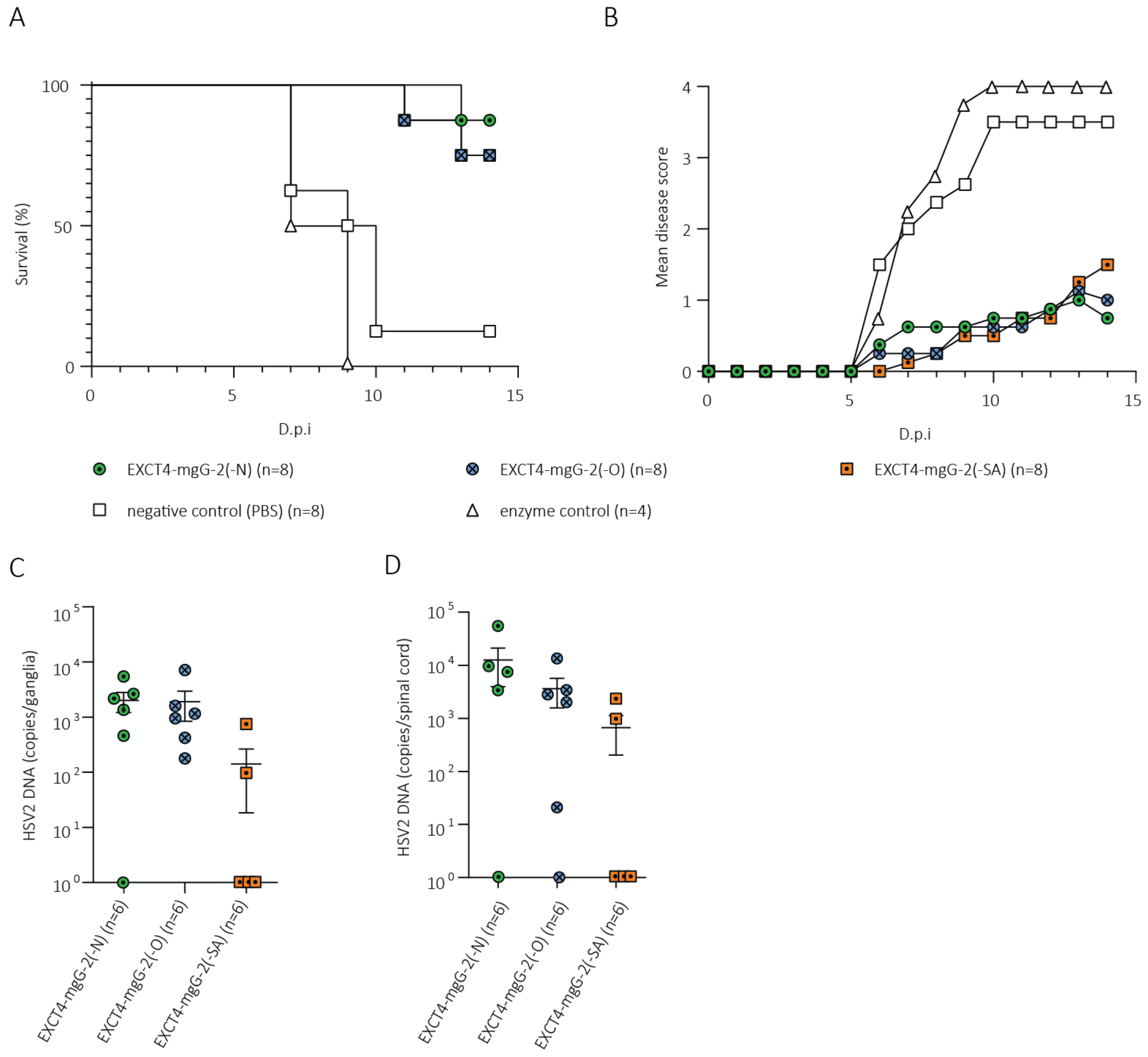


**Figure S3.** C57BL/6 mice were intramuscularly immunized with EXCT4-mgG-2(-N), EXCT4-mgG-2(-O) and EXCT4-mgG-2(SA) and genitally challenged with 25 x $LD_{50}$ of HSV-$2_{WT}$. The survival rate **(A)** and disease score **(B)** was assessed until 15 d.p.i. Viral spread to neuronal tissue; HSV-2 DNA copies per ganglia **(C)** HSV-2 DNA copies per spinal cord **(D)**. Statistical analysis was performed with the pairwise log-rank (Mantel Cox) (A) or Kruskal-Wallis test (C-D). The detection limit for HSV-2 DNA in ganglia and spinal cord was 40 and 160 copies respectively. D.p.i = Days past infection. Values are expressed as means ± SEM.

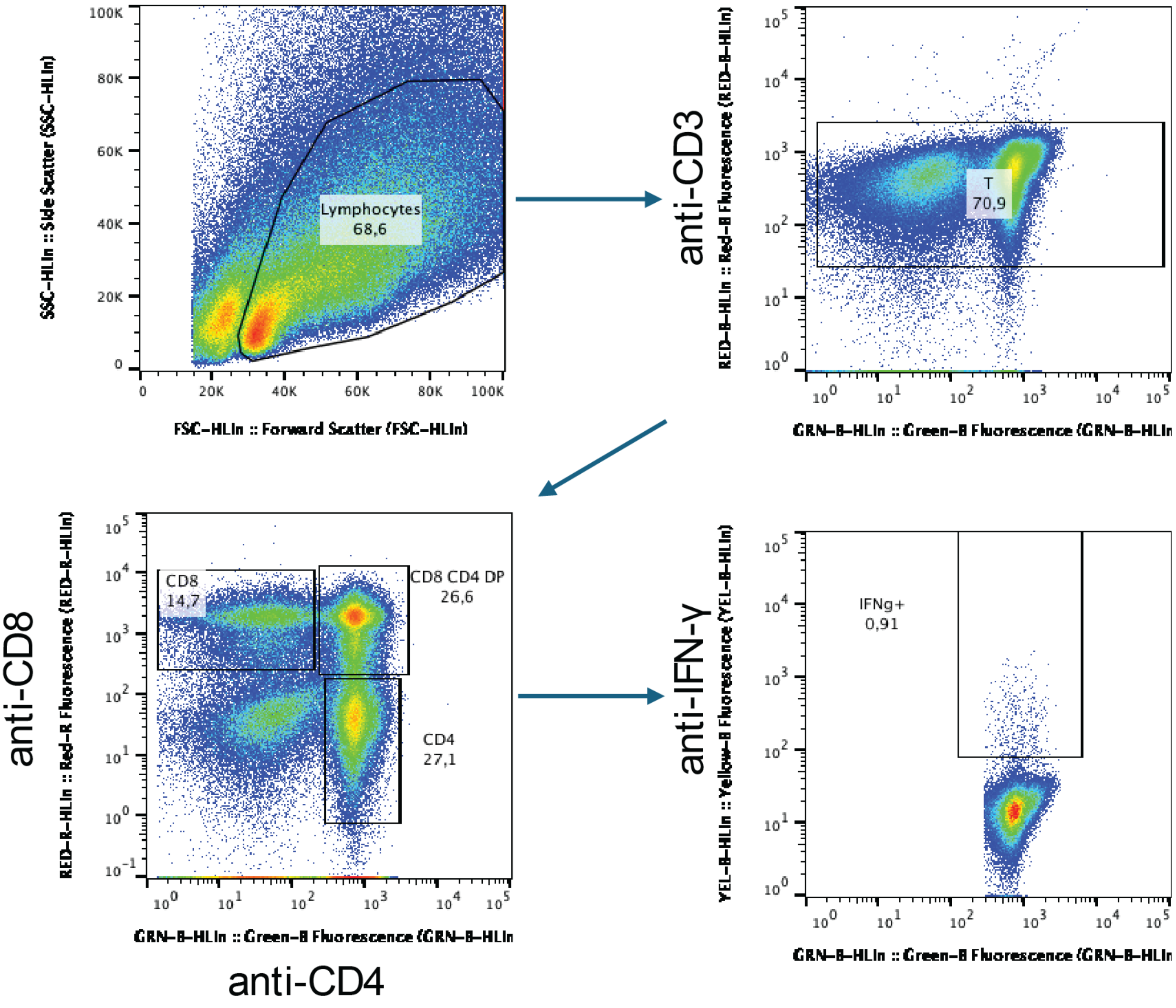


**Figure S4**. Representative flow cytometric analysis of intracellular INF-γ stained CD4+ T cells. Outlined is the gating strategy used for both CD4+ and CD8+ T cells.

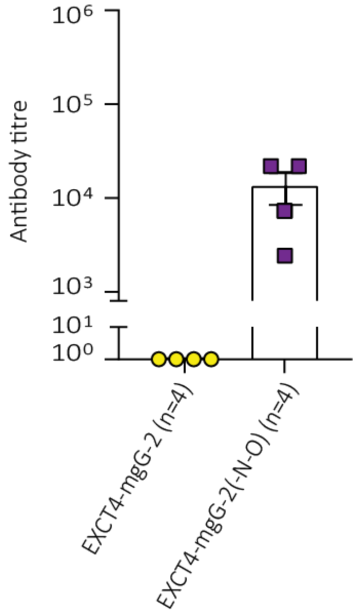


**Figure S5.** Serum antibody reactivity against the glycosidases present in the EXCT4-mgG-2(−N−O) preparation. Serum samples was collected 14 days following immunization with a preparation containing glycosidases, but no recombinant mgG-2, and the reactivity against the EXCT4-mgG-2 and the EXCT4-mgG-2(−N−O) assessed. Presented is mean ± SEM.

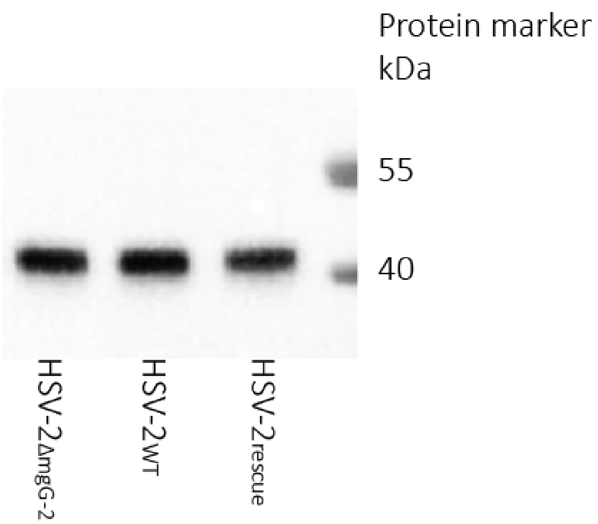


**Figure S6.** Western blot of cell lysates and growth medium after infection of Hep-2 cells with HSV-$2_{WT}$, HSV-$2_{\Delta mgG\text{-}2}$ or HSV-$2_{rescue}$. Anti-sgG-2 mAb recognized sgG-2 (44 kDa). One experiment of two performed is shown.
